# Root hair growth phases are coordinated by cytoskeleton, nucleus dynamics and cell mechanics in Arabidopsis

**DOI:** 10.1101/2025.03.31.646268

**Authors:** Gilles Dupouy, Tamsin Spelman, Gaurav Singh, Etienne Herzog, Stephanie Baudrey, Simone Bovio, Jérome Mutterer, Olivier Hamant, Alexandre Berr, Henrik Jönsson, Marie-Edith Chabouté

## Abstract

Polar cell growth is a fundamental process across organisms, yet its coordination with nuclear movement and cytoskeleton dynamics remains underexplored. Focusing on *Arabidopsis thaliana* root hairs, we investigate these processes using high-resolution live imaging within microfluidics-based experiments. By incorporating data on cytoskeletal dynamics, nuclear positioning, and tip growth into a mathematical model, we analyse how their interactions shape the different growth phases that we reveal for the first time in this study. Chemical treatments and mutant analyses further support our model, revealing that timely cytoskeletal changes drive transitions between these growth phases, and correlate with shifts in nuclear movement and morphology. This regulation suggests a microtubule-actin crosstalk in the root hair subapical region. Additionally, we present novel findings on vacuole movement and cell stiffness, further refining our understanding of tip growth dynamics. Collectively, our work provides a comprehensive framework for understanding how transitions between growth phases are orchestrated in plant tip-growing cells.

## Introduction

Over the past few decades, models of polar cell growth have been developed using tip-growing cells across various systems, including neural axons in mammals (*1*) filamentous tissues in moss (*2*), bacteria (*3*) plant zygote elongation (*4*) or specialized plant cells like pollen tubes and root hairs (RHs) (*5*). RHs are rapidly growing tubular extensions of root epidermal cells, essential for plant water and nutrient uptake, directly or indirectly through establishing symbiosis such as with mycorrhizal fungi (*6, 7*). RH must regulate its transition from initiation via rapid elongation to tip growth arrest, while ensuring cylindrical shape maintenance through precise cell wall synthesis(*8, 9*).

In growing RHs (GRHs) with rapid elongation, both cortical microtubules (CMTs) and highly dynamic endoplasmic microtubules (EMTs) coexist in the shank, i.e. the zone of secondary cell wall deposition at approximately 20µm from the tip in average (*10*) with MTs nucleating on the nuclear surface and cortex (*11, 12*). Actin filaments (AFs) form bundles in the shank, while dynamic fine AFs localize with EMTs in the subapical region (∼5µm from the tip), with globular (G)-actin concentrated at the tip (*12–14*). Actin polymerization drives polar growth in Arabidopsis RH cells (*15*). As RHs become mature (MRHs), fine AFs and G-actin at the tip are replaced by thick AF bundles (*13*), and EMTs disappear, leaving CMTs looping around the tip and body (*12*). MTs regulate cell shape, directional growth (*16*), and cellulose deposition (*17*), while AFs support intracellular transport critical for tip extension (*18*).

During rapid tip elongation (ranging from 1 to 2.5 µm/min), the nucleus maintains a nearly constant distance from the tip (ranking between 60 and 80 µm) crucial for tip growth (*19*). Such nucleus connection to the tip is lost in MRHs and nuclear movement is then solely regulated by AFs (*19, 20*). Nuclear movement is not purely unidirectional in growing RHs. Instead, a small consistent back-and-forth motion has been observed, which may be controlled by AFs and was shown to be dampened by MTs (*21*). A correlation between tip growth rate (TGR) and nuclear position has been noted in response to increased substrate rigidity, though the underlying mechanisms remain unknown (*22*). Throughout RH development, nuclear shape evolves, with its aspect ratio (AR), i.e. the ratio of its longest length divided by its smallest, increasing progressively as the cell matures (*20, 23*). This elongation is influenced by mutations affecting the nuclear lamina, LINC complexes (Linker of Nucleoskeleton and Cytoskeleton), Myosin XI-I , and several nucleoporins (*24, 25*).

Several computational models have been developed to explain specific aspects of RH growth and structure. Simpler mathematical models have addressed cell wall expansion using scaling arguments (*26*), while more complex models incorporate biomechanical frameworks (*27*) and cell wall aging dynamics (*28*). Cytoskeletal features have also been investigated, with MT alignment modelled using mean-field theory (*29–31*) and computational simulations (*32–34*). However, AF alignment in plant cells remains less explored in models. Most cytoskeletal network models consider MTs and AFs independently of RH development, though some address cytoskeletal force distribution, such as local stresses on MTs during RH initiation (*35*). Computational studies on nuclear deformation in RHs and plant cells are limited, despite broader research in other cell types (*36, 37*).

Tip growth primarily occurs at the apex, with limited length extension in the shank (20µm from the tip), through the synthesis of a primary cell wall (*9*). Meanwhile, shank hardening is facilitated by the deposition of a secondary cell wall layer (*8*). Both cell wall thickness and composition, as well as the underlying turgor pressure contribute to overall RH stiffness. Atomic force microscopy (AFM) analysis revealed a reduced cell stiffness in the shank of mutants with impaired CMT organisation (*38*). The central RH vacuole plays a key role in maintaining turgor pressure (*39*) and expands progressively during RH growth (*40*). Initially confined to the basal shank, it extends into the subapical region and reaches the tip in MRHs (*41*).

While various cellular parameters—such as cell wall stiffness, cytoskeletal organization, vacuole and nuclear dynamics— have been mostly investigated independently in RH growth, an integrated framework linking these factors remains lacking. To date, no theoretical model has fully integrated the mechanistic interactions between the cytoskeleton, tip growth, and nuclear movement and shape. Building on these gaps, this study aims to decipher the mechanistic of tip growth during *Arabidopsis thaliana* RH development through the building of an integrative model focusing on key cellular components. Specifically, we seek to identify critical behaviours that regulate the timely progression from the growth phase to tip growth arrest. Leveraging our recently developed high-resolution imaging technology combined with microfluidics (*23*), we quantitatively analyse TGR, nuclear dynamics (movement and shape), and cytoskeletal organisation from rapid tip elongation to tip growth arrest. Our findings reveal a key developmental transition in growth dynamics between rapid growth (G) and reduced growth stages (RG), governed by cytoskeleton-related TGR shifts to assure the timely maturation of RHs. Using these data, we quantitatively characterise cytoskeletal changes during each of these transitions in relation to nuclear positioning and tip behaviour. The development of a mathematical model enables the investigation of regulatory mechanisms connecting modifications of nuclear motion and morphodynamics during tip growth based on cytoskeleton and nucleus generated forces. To validate our model, we apply cytoskeleton-destabilizing drugs and analyse mutant lines affecting MT arrays (*fra2*) and nuclear morphodynamics (*crwn1-2*). Our findings present a comprehensive model linking cellular tip growth to nuclear positioning *via* the cytoskeleton. The dynamics following chemical perturbations suggest that this G/RG transition depends on major changes in cytoskeleton dynamics, with decreased MT stability but increased AF stability. We showed a forward shift of the nuclear position during this transition, which is mirrored by a similar shift in vacuole position, suggesting an additional role for the vacuole movement in this process. This also results in local changes in cell stiffness at the subapical region, highlighting this particular region for the control of RH growth. Following the G/RG transition we identified a timely controlled decrease of TGR until arrest, correlated with a cytoskeleton and nucleoskeleton-controlled stretching of the nucleus.

## Results

### Cytoskeletal reorganisation and nucleus repositioning correlate with transitions between root hair growth phases

Cytoskeleton networks undergo organizational changes during RH development, but quantitative analysis, particularly regarding TGR and nucleus position, is lacking. To address this, we collected consistent experimental data using our previously described microfluidic setup (*23*). Our study spans RH development from growth to early maturation after tip growth arrest. Real-time confocal imaging in microfluidic channels (*42*) was performed every 3 mins on 9-day-old *Arabidopsis* seedlings. Using differential interference contrast (DIC), we measured TGR kinetics until growth arrest (Fig. 1A). Initially, RH growth remained relatively constant at a mean rate of 1.1 ± 0.2 µm/min, then switched to a reduced growth phase characterized by a gradual decline of TGR until arrest, with a mean rate of 0.29±0.04 µm/min (Fig. 1B, n_G_=7, n_RG_=11, p=0.003, Mann Whitney test). This pattern allowed us to define three RH development stages: the Growth (G) stage, lasting up to 150 min before tip growth arrest, with a TGR ranging from 0.6 to 2 µm/min; the Reduced Growth (RG) stage, defined by TGR < 0.6 µm/min; and the Early Mature (EM) stage beginning after tip growth arrest (Fig. 1A). A clear transition was detected between the G to RG phases, (hereafter G/RG) (Fig. 1C, fig. S1D), characterized by a sharp 35% TGR drop occurring within 20 min (Fig. 1D, from 0.9±0.1 to 0.6±0.1 µm/min, n=9, p=0.04).

**Fig. 1:**
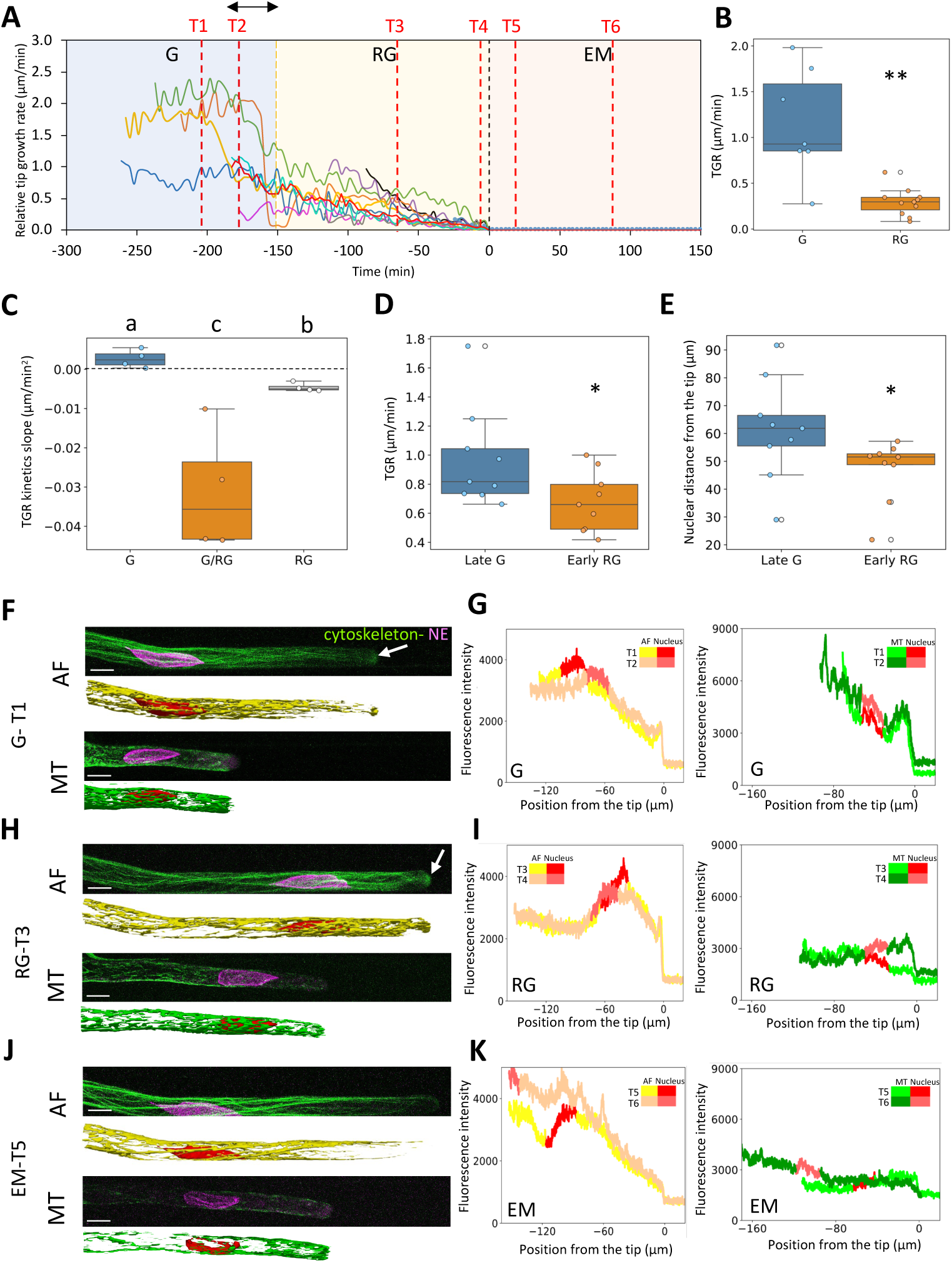
Kinetics of RH tip growth linked to cytoskeleton reorganization. **(A)** Evolution of Tip Growth Rate (TGR) over the three development phases: Growing (G), reduced growth (RG), and early maturation (EM). Each line represents individual RHs (n=11). Black double arrow represents the transition between G and RG phases. (**B**) TGR mean values in G and RG stages, ** P-value < 0.01. (**C**) TGR kinetics slopes for G and RG phases, as well as the G/RG transition in between (black double arrow on A). Different letters highlight statistically different conditions according to Mann-Whitney non parametric test (p < 0.05). (**D**) Mean TGR at the end of G phase and beginning of RG phase (around G/RG transition). (**E**) Nuclear distance from the tip at the end of G phase and beginning of RG phase (mean values from 10-minute timeframes right before and after G/RG transition). * P-value < 0.05. (**F** to **K**) Confocal microscopy images and cumulated Microtubule (MT) and Actin filament (AF) fluorescence intensity of RHs from each phase: G (**F** and **G**), RG (**H** and **I**), and EM (**J** and **K**). White scale bar: 10µm. AF (yellow) and MT (green) intensity profiles corresponding to the images from F (**G**), H (**I**) and I (**K)** respectively. On each graph two timepoints are represented, corresponding to the T1 to T6 time points displayed on A. the nucleus position is highlighted in red on each curve. White arrows indicate the presence of G-actin.

Next, we analysed AF, MT and nuclear organisation changes across RH growth kinetics in seedlings co-expressing SUN2-tagRFP with either GFP-MBD (for visualizing MTs) or LifeAct-GFP (for AFs). During the G phase, the MT bundling peak was located at the back of the nucleus, with a smaller peak at the front (15–5 µm from the tip, *i.e.* the subapical region) corresponding to the MT fringe, coexisting with fine AFs while the diffuse fluorescence signal at the tip (Fig. 1F, white arrows) may correspond to fine AFs and eventually G-actin as previously reported (*14*). Further upstream (∼20 µm from the tip), both MT and AF bundles were present in the shank. At G/RG (n_G_=16), AF intensity shifted towards the subapical region, while MT intensity decreased (Fig. 1H, I, fig. S1B), coinciding with a rapid cytoskeletal reorganisation (movies S1, S2). These organisation patterns remained stable through the RG phase. Just before tip growth arrest, a transient MT bundling occurred at the tip (∼20 min, movie S3), while AFs polymerized into longer filaments (movie S4). After tip growth arrest, AF intensity peaked further back (after T5), while MT intensity remained low (Fig. 1J, K), reflecting EMTs disappearance and the decreased density of CMTs as previously reported (*12*).

Nuclear position correlated with cytoskeletal changes. During the G phase, the nucleus was 61±6µm (Fig. 1E) from the tip, aligning with previous observations (∼60–80 µm (*19*)). AF intensity peaked around the nucleus, while MT intensity peaked slightly behind it (Fig. 1G, 69% of RHs, n_G_=13). At G/RG, the nucleus moved closer to the tip, reducing the gap to 47 ± 3 µm (Fig. 1E, n_G,_n_RG_=9, p=0.03 compared to G phase), which coincided with the forward shift of the AF intensity peak (Fig. 1H, I). The nucleus position then remained stable through RG (movies S1 and S2). After the beginning of the EM phase, the nucleus shifted backward, mirroring AF intensity retreat (Fig. 1J, K, movie S5). Fully mature RHs showed more variation in cytoskeletal density distribution between the nucleus and the tip in comparison to G and RG phases (fig. S1C).

In summary, cytoskeletal reorganisation closely aligns with nuclear movement, first advancing toward the tip at the G/RG transition and later retracting in EM phase.

### Mathematical model of RH growth coupled to nucleus dynamics

To move beyond correlations between cytoskeleton, nucleus, and tip growth, and try to decipher the causal effects between these elements, we developed a mathematical model based on the different stages of RH development (Fig. 2, Methods, text S1). We adapted a previous mechanical model for tip-growing cells (*27*), which incorporates plant cell specific features such as viscoelastic cell wall properties and turgor pressure. We extended this model by coupling tip growth with nucleus dynamics through cytoskeletal forces (Fig. 2A, fig. S2A, text S1). We simulated the RH as rotationally symmetric about its long axis, resulting in a generally cylindrical cell shape (but allowing for variation in RH diameter) which smoothly transitions into an evolving rounded cap in the tip growth region. To model the evolution of TGR, we adjusted the extensibility magnitude of the tip in simulations to match the experimentally observed growth rates across the three growth phases simplifying the rate progression into a continuous, piecewise linear function (Fig. 2B). The nucleus, modelled as a constant-volume spheroid, is tethered to the tip by two spring forces representing the net force generated by MT and AF. According to the changes in cytoskeleton patterns observed from growth to RH maturation (Fig. 1), we adjusted the AF and MT force potentials relative to the nucleus position from G phase through to EM phase (fig. S2C). Based on prior findings (*21*), we assumed a prevalence of AF over MT effects on nuclear movement. An additional spring force from the nucleoskeleton accounts for nuclear envelope tension, allowing for nuclear shape changes, from squeezed to elongated, influenced also by both cytoskeletal forces. By adjusting model parameters, we accounted for changes in tip extensibility and cytoskeletal forces during RH development. A crosstalk between the AF and MT networks was also incorporated (text S1). We simulated the growth of a RH initiating from a trichoblast cell over 600 min and into early maturation after tip growth arrest (text S1). For simplicity, we started with a spherical nucleus in the centre of a spherical cell representing a trichoblast cell (Fig. 2C). Other trichoblast cell shapes can also be used but had no significant effect on our results. As the simulation progressed, the long thin RH protrusion grew at an approximately constant rate. When the cell was 72 µm long (around 20 min post initiation, movie S6), the cytoskeleton forces on the nucleus turned on and the nucleus moved into the protrusion and started following the tip at a constant average distance with small fluctuations of its position from -520 to -200 min (Fig. 2C-D, fig. S2B-C). In this growing phase (G phase), based on observed cytoskeleton peak intensities (MT peak behind the nucleus and AF peak at the nucleus position, Fig. 1F-G), we positioned the AF and MT force potentials respectively so that AF forces pull the nucleus towards the tip while MT forces preferentially provide a force pulling the nucleus away from it (fig. S2C). Therefore, the nucleus remained close to but behind the equilibrium position of AF forces and at the front of the equilibrium position of MT forces, and this equilibrium position followed the tip growth. After 400 min of growth, we prescribed the start of G/RG by reducing tip extensibility over 50 min, causing TGR to be reduced (Fig. 2B red line, fig. S2B, D). This was followed by a 150 min time frame representing the RG phase with tip extensibility linearly reducing to zero. After an initial reduction in MT forces during G/RG, these forces were reduced linearly to 0 through the RG phase to reflect the large decrease in MT intensity seen in experiments (Fig. 1). Simultaneously, the nucleus moved forward with its position becoming solely determined by the centre of AF forces (Fig. 2D). After 600 min, tip growth stopped and we prescribed a retreat of the centre of AF forces leading the nucleus to move away from the tip during the EM phase (Fig. 2C-D, red line in D, fig. S2B-C). From growth to maturation, our model predicted the nucleus to change morphologically (Fig. 2G,I, red line). During the G phase, the nucleus was pulled from the back towards the tip, causing it to become compressed and resulting in a low aspect ratio (AR) and area. At the G/RG transition, the nucleus moved forward, reducing this squeezing effect leading to an increased nuclear AR. This however also resulted in the progressive increase of the nucleus area (Fig. 2I, J). Finally in the EM phase, the retreating AF effect resulted in a slight elongation of the nucleus. The AR changes were inversely correlated with circularity (fig. S3A) and the mean nuclear solidity remained unchanged in our mathematical model as the nucleus was assumed spheroidal (fig. S3C). A model parameter sensitivity analysis identified the importance of individual parameter values to the measured behaviours (fig. S2A, text S1).

**Fig. 2:**
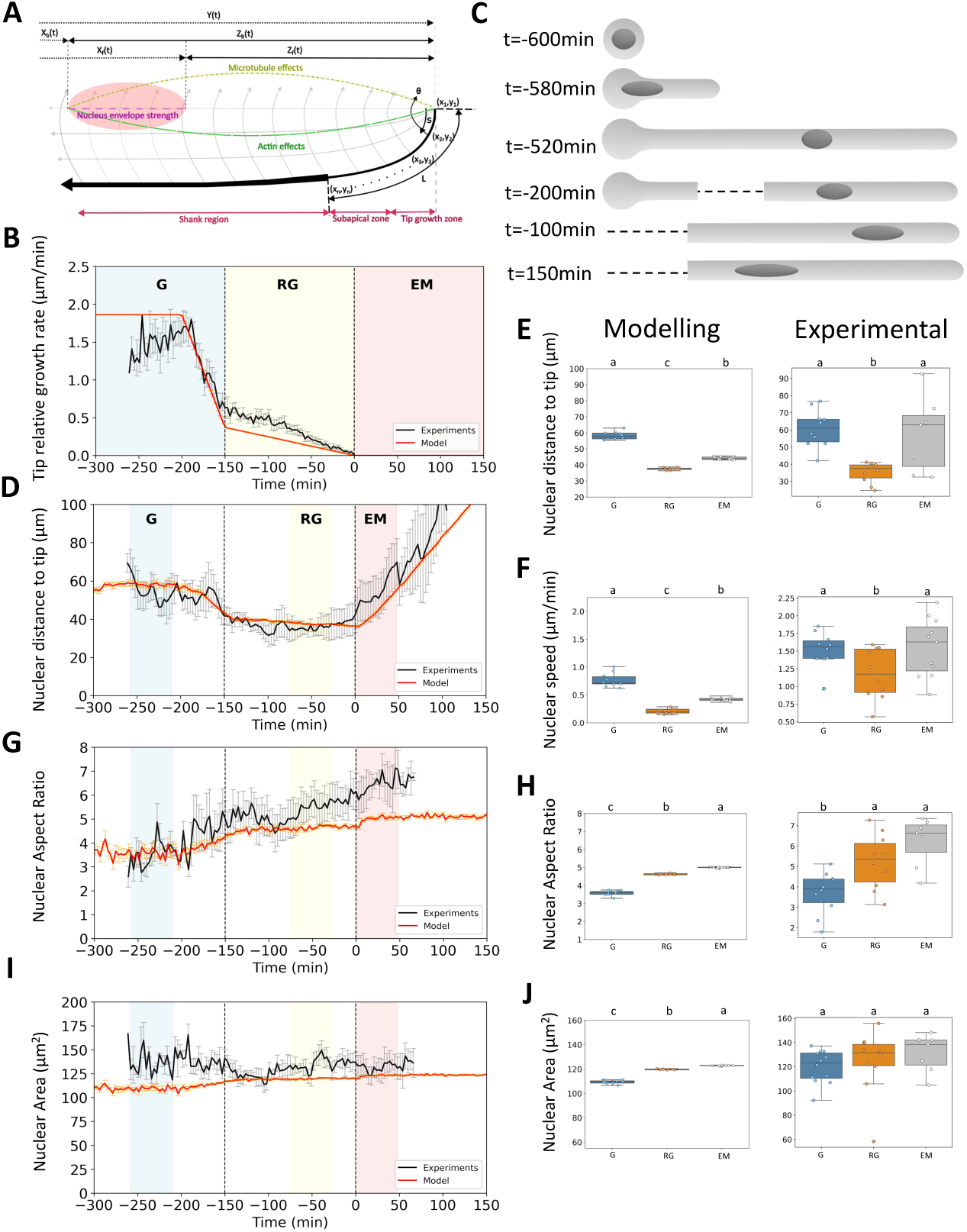
RH growth and nuclear dynamics. **(A)** Modelling of RH tip growth with cytoskeleton and nuclear dynamics. (**B**) Superposition of RH TGR experimental (black line) and model (red line) kinetics. (**C**) Morphological drawing of RH tip elongation and nuclear movements according to the model proposed in A. (**D**, **G** and **I**) Superposition of RH experimental kinetics (black line) and model rendering data for nuclear distance to the tip (D), nuclear aspect ratio (G) and nuclear area (I). Blue, yellow and red boxes highlight the time frame used to produce average values for each growth phase. (**E**, **F**, **H** and **J**) Average values of model (left) and experimental (right) data for nuclear distance to the tip (E), nuclear speed (F), nuclear aspect ratio (H) and nuclear area (J) between each growth phases. Different letters highlight statistically different conditions according to Mann-Whitney non parametric test (p < 0.05).

Thus, building on a mathematical model of a tip growing cell developed by Dumais et al. (*27*), we specialised it for RH cells and expanded it by coupling tip growth to nuclear dynamics (movement and shape) via cytoskeleton forces to model RH growth until maturation. In the next section, we experimentally tested the predictions of this model.

### Accuracy of model predictions compared to experimental data

To test the accuracy of our model, we compared our simulations with experimental data. We have simulated tip growth and nuclear dynamics from growth to maturation and compared TGR and nuclear parameters over a 48-min window for each phase (G, RG and EM). In the G phase, TGR and nuclear positions are aligned with experimental data (Fig. 2B-E) with a mean TGR of 1.5±0.1 µm/min and nuclear position of 60±3 µm/ tip (n=10). Then in the RG phase, the model assumed a 2-fold decrease of TGR (Fig. 2B) which led to a predicted change in nuclear position relative to the tip as observed experimentally (Fig. 2D-E). Retreat of the nucleus after tip growth arrest in the EM phase indicated a loss of its connection to the tip, as seen in our data (Fig. 2C-E). In simulations, nuclear speed followed the TGR decrease and the nucleus forward movement from G to RG phases, but remained relatively low in EM compared to our experimental values although being higher than in RG (Fig. 2F, fig. S2E). Altogether, the model is able to accurately predict the nuclear movement during the transitions between growth phases.

Concerning nuclear morphodynamics, simulations predicted an increase of nuclear AR in the RG phase as observed experimentally (Fig. 2G, H). However, no significant difference was observed between RG and EM phase experimentally, despite the kinetics apparently not changing from the RG phase, which is probably due to the variability observed between individual RHs. Simultaneously the nuclear circularity statistically decreased both in the mathematical model and experimentally (fig. S3A, B). Similarly, whereas our simulations predicted a slight increase in the nuclear area in RG phase, the experimentally measured maximum-projected nuclear area did not show any significant change through the whole maturation kinetics (Fig. 2I, J). Finally, the unchanged nuclear solidity in the model matched the statistically unchanged solidity from experimental data (fig. S3C, D). Besides the squeezing effects imposed in our model by cytoskeleton forces we cannot exclude a role of the nucleoskeleton to counterbalance these effects. For example, CRWN1, one of the nucleoskeleton components, was shown to control nuclear expansion (*43*).

In summary, our mathematical model is accurately able to capture changes in TGR with nuclear movement behaviour observed experimentally, suggesting that this model has a good predictive power. Next, we tested whether predictions from perturbations within the model could be observed experimentally, notably through cytoskeleton destabilisation.

### MT dynamics regulates G/RG transition and tip-nucleus connection

We observed correlations between cytoskeleton organisation, nuclear position and TGR. Our model further suggests a causal link between these events, indicating that the observed decrease in TGR coinciding with the decrease in MT intensity at the G/RG transition may be sufficient to explain the reduced nuclear position relative to the tip. To explore this possibility, we first tested the effects of a MT destabilisation on TGR and nuclear position during the G phase, since a destabilisation of AF is known to lead to tip growth arrest (*13, 16*). In our model, we simulated this destabilisation by reducing the MT force on the nucleus to zero and then partially recovering it to simulate drug removal. We also incorporated cell extensibility changes to modify tip growth speed. For both model and experiments we have used the same pipeline of analysis. After a 10-min control in G phase, we induced a cytoskeleton depolymerisation for 10 min and tracked RH behaviour for 90 min post-drug removal. Experimentally, we induced a MT destabilisation using 1 µM Oryzalin (OZ) and used the same fluorescent reporting lines SUN2-tagRFP with either GFP-MBD (for MTs) or LifeAct-GFP (for AFs) to evaluate a potential AF-MT crosstalk (fig. S4). Live imaging was conducted for 110 min over the whole experimental framework. Mock controls were performed under identical conditions using liquid growth medium.

In simulations, the reduction in TGR was imposed by a decrease in tip extensibility, partially recovered upon repolymerisation of MTs which interplays with AFs (Fig. 3A model, text S1). In our experiments MT depolymerisation caused AF bundle formation in the subapical region which was further maintained during recovery (fig. S4B, see white arrow), correlating with a significant decrease in TGR (Fig. 3A, p=0.04, n=10). This suggests that the effect of MT depolymerisation affects TGR through tip extensibility via MT-AF crosstalk. This could be linked to vesicle trafficking which was previously shown to be regulated by AF dynamics (*44*). In the recovery stage, a partial recovery of the TGR was stabilized at a low level, mimicking a RG growth phase, which was imposed in the model (Fig. 3A). This relies experimentally on a very slow repolymerisation of MTs occurring in the shank and the subapical region while AF bundles were even more increased (fig. S4B, see white arrow). A simulation of MT destabilisation had no effect on nuclear distance from the tip during treatment and early recovery, with a backward movement of the nucleus from mid-recovery onwards (Fig. 3B). This was explained by AF-MT crosstalk resulting in a weakening of the AF link to the tip causing the nucleus to not move as fast as the tip during recovery (fig. S4A). Experimental data showed a similar trend in terms of kinetics (Fig. 3B, C), however with already a significant backwards movement in the early recovery (ER) phase, further paused upon MT repolymerisation (Fig. 3B, D). As the MT repolymerisation was not complete, this backward movement likely reflected AF forces on the nucleus with MT forces gradually counteracting this effect, suggesting a more direct connection of the nucleus to the tip via MTs and a backward force of AFs. Nuclear speed was predicted to remain unchanged in the model, aligning with experimental data (Fig. 3E, n=6).

**Fig. 3:**
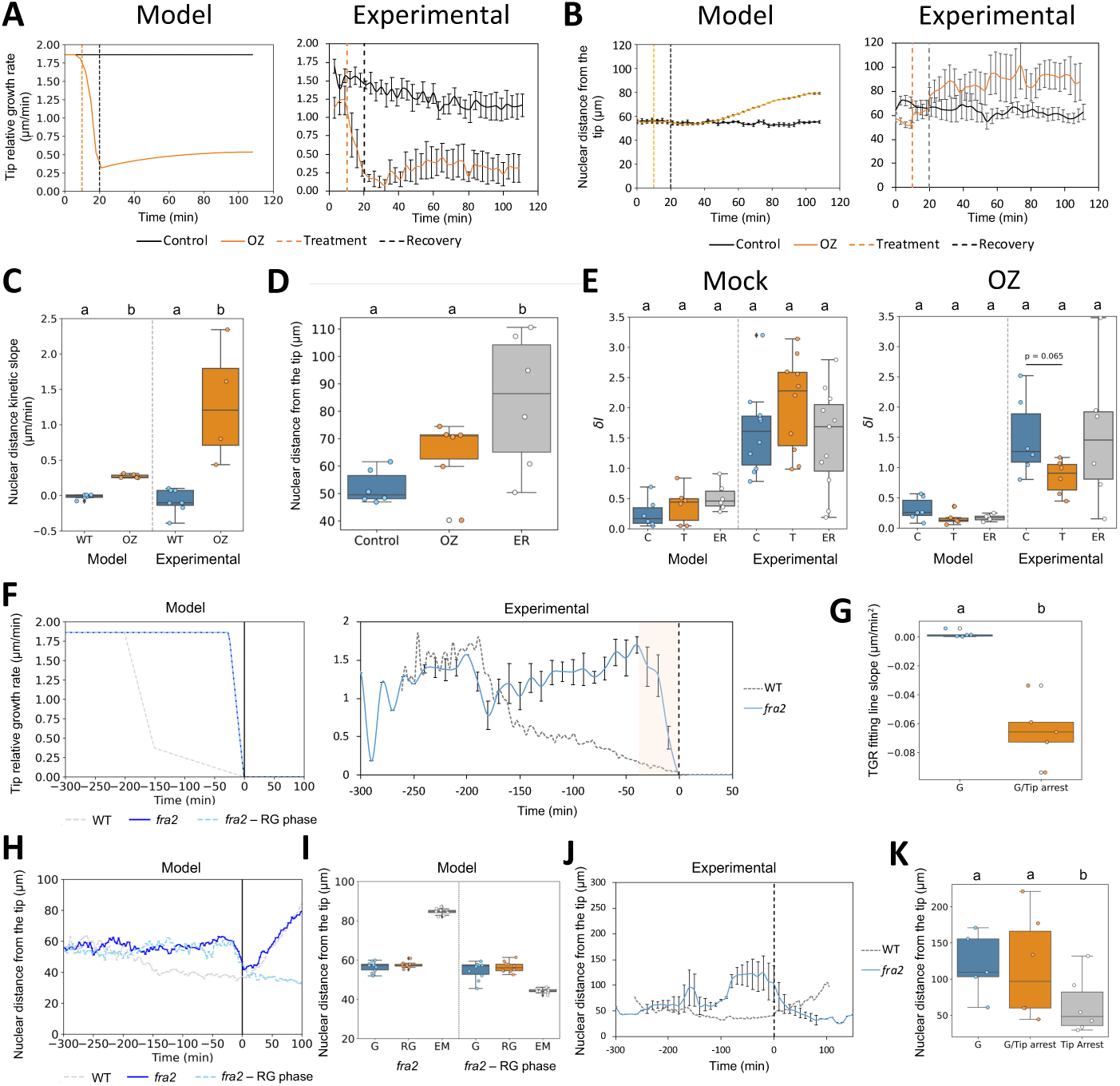
A transient destabilization of MTs triggers the transition from G to RG phases. **(A)** Kinetics of TGR of G phase RH in mock conditions (black line) or with 1µM Oryzalin (OZ) (orange line) from simulations (left) and experimental data (right). (**B**) The kinetics correspond to a 10 min-control medium, followed by a 10 min-treatment, and a 90 min-drug removal. (**B**) Kinetics of nuclear distance from the tip in G phase in mock conditions (black line) or with 1µM Oryzalin (OZ) from simulations (left) and experimental data (right). (**C**) Kinetic slope of nuclear distance from the tip for Control (WT) and OZ treatment (OZ) from simulation (left) and experimental data (right). (**D**) Nuclear distance from the tip in G phase control, OZ treatment and early recovery (ER). (**E**) Nuclear speed in G phase control, treatment (mock on the left, OZ on the right) and recovery from simulation (left half panel) and experimental data (right half panel). (**F**) TGR kinetics observed for *fra2* mutant (blue lines) compared to WT Col-0 (dashed grey line) from simulation (left) and experimental data (right). The orange box highlights the transition between G and tip growth arrest experimentally. In simulations, we considered 2 scenarios for *fra2* mutant: either i) absence (blue line) or ii) presence (dashed blue line) of a RG-like phase following tip growth arrest. (**G**) Fitting line slope for G phase and transition into tip arrest in *fra2* mutant. (**H**) Simulations of nuclear distance from the tip in *fra2* mutant with the 2 scenarios described in (F). (**I**) Average nuclear distance from the tip at each growth phases from model data according to the 2 scenarios described in (F) (without RG, left and with RG, right) (**J**) Kinetics of nuclear distance from the tip for *fra2* mutant (blue line) compared to WT Col-0 (dashed line) from experimental data. (**K**) Average nuclear distance from the tip at each growth phases in *fra2* mutant (G, G/RG-like transition and Tip arrest). Different letters highlight statistically different conditions according to Mann-Whitney non parametric test (p < 0.05).

To explore the effect of MT stabilisation on TGR and nuclear movement, we used a 10µM Taxol treatment. This treatment had no significant effect on TGR (fig. S5A) or nuclear position (fig. S5C), consistent with the fact that a relative MT stabilisation already exists in G phase RHs and may be important for maintaining the nuclear position relative to the tip. This could be linked to the observed EMTs bundling in the subapical region (Fig. 1) with concomitant AF bundling there (fig. S4C, white arrows) which was maintained upon MT stabilisation, probably preventing the nucleus from moving forward. To further investigate the regulation of MT stability, we examined the *KATANIN1* mutant *fra2*, which displays a reduced MT severing activity affecting MT arrays and resulting in longer RHs (*45*). The *fra2* mutant showed an enlarged MT zone around a more rounded nucleus (fig. S4E), and maintained a high TGR resembling the G phase from WT lines (1–1.8 µm/min, n=5, Fig. 3F). In simulations, we adjusted the tip extensibility to match the experimental TGR pattern while maintaining a high MT strength throughout (text S1). Our model tested two scenarios: i) either the tip growth arrest in *fra2* aligns with the start of the EM phase (normal WT tip growth arrest) or ii) the tip growth reduction corresponds with the G/RG transition. Our simulations show that in the second scenario, the nuclear movement is more consistent with our experimental data (Fig. 3H-K), with the TGR quickly dropping within 25–50 min before tip growth arrest and the nucleus not retreating within the 150 min post tip arrest (Fig. 3J). Statistical analysis of TGR slopes confirmed a significant drop of TGR before tip growth arrest which was not observed in WT (Fig. 3G). It overall suggests that in this mutant, even though the tip arrests prematurely compared to WT (before any RG phase can be triggered), the nuclear behaviour appears to follow the WT course of events for RH maturation (G -> RG -> EM) and maintains this G/RG transition. Alternatively, stabilising AFs with 5 µM Jasplakinolide (JP) led to tip growth arrest (fig S5A), possibly relying on its described role for G-actin polymerization (*46*) at the tip which will halt vesicular trafficking (fig. S4D).

In summary, we show experimentally that both MT destabilisation and AF stabilisation trigger a G/RG-like effect on TGR. In addition, while a complete MT destabilisation also results in reduced TGR, it also leads to a loss of coordination between TGR and nuclear position. The model suggests a possibility to explain this scenario that is to have a cross-talk between MTs and AFs (fig. S4A), consistent with the increased AF bundling in the subapical region in OZ treatments. On the other hand a MT over-stabilisation maintains the G/RG transition effect on nuclear movement, but results in a coinciding tip growth arrest. Consequently, MT stability needs to decrease to trigger G/RG transition while remaining within a defined range for the nucleus-tip connection to be maintained.

### Timely cytoskeleton stabilisation is needed for tip growth arrest

We predicted changes in MT and AF effects for RH maturation (Fig. 2). Therefore, we tested cytoskeleton destabilisation during RH maturation, simulating MT or AF destabilisation in late-RG phase RHs using the same analysis pipeline as above.

Similarly imposed decrease of tip extensibility (text S1) in simulations caused a TGR decline upon MT destabilisation by OZ (Fig. 4A, model OZ), which was partially recovered within 90 min post treatment. This relies experimentally on an increase of AF bundling, especially in the subapical region that was maintained upon recovery (Fig. 4A, n=6, p=0.02, fig. S6A, see white arrows). This suggests that AF bundling is key for controlling tip growth as we observed for RH maturation (Fig. 1). However, while simulations showed no change in the nuclear position during treatment and recovery, we observed a mean backward movement of the nucleus from early to late recovery (Fig. 4B). This may rely on the observed decrease in AF fluorescence intensity at the front of the nucleus during recovery (fig. S6A) allowing this retreat of the nucleus which was not considered in our simulation. Upon AF destabilisation, we imposed a reduction of tip extensibility to zero which resulted in no change in the nucleus position in the model as observed in our experiments (Fig. 4B). Of course, a similar effect was predicted and observed upon a destabilisation of AFs during the G phase (fig. S7 A-B) as previously described (*13*). Contrary to the scenario observed in G phase (fig. S7A), the destabilisation of AFs in the RG phase could not be completed in the subapical region after 10 min of LB treatment, suggesting the existence of a more stable AF network in this phase compared to G phase (fig. S6B, white arrow). Additionally, AF destabilisation led to MT destabilisation post-drug removal, highlighting AF-MT crosstalk (fig. S6B). In support of this, the destabilisation of both networks was delayed upon a cumulated OZ and LB treatment and their repolymerisation was very slow upon recovery (fig. S6C).

**Fig. 4:**
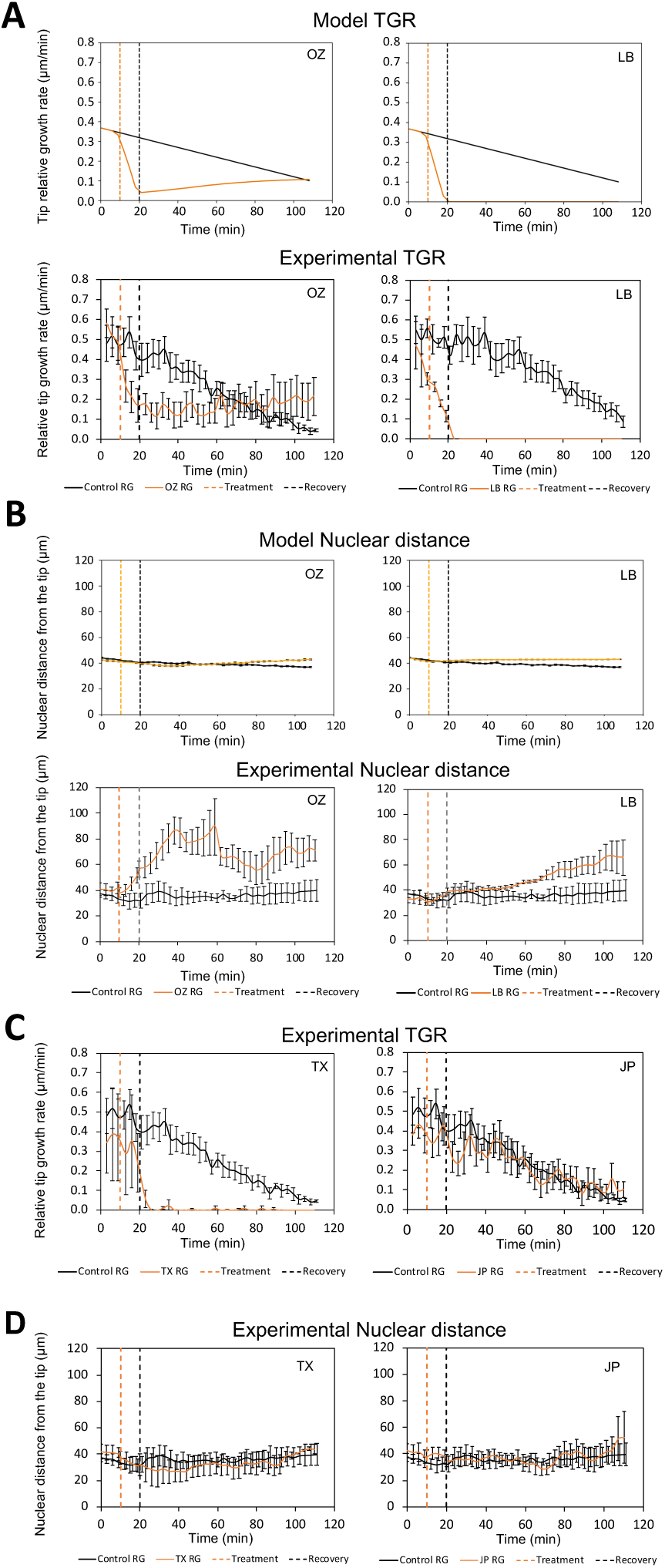
Crosstalk between MT and AF is necessary for timely RH maturation. (**A**) Kinetics of TGR of RG phase RHs in mock conditions (black line) or treated (orange line) either with 1µM Oryzalin (OZ) or 1µM Latrunculin B (LB) from simulations (above) and experimental data (below). The kinetic of treated RHs (orange line) corresponds to a 10-min control medium, followed by a 10 min-treatment, and 90 min-drug removal. **(B)** Kinetics of nuclear distance from the tip of RG phase RHs in mock conditions (black line) or treatments (orange line) with either 1µM Oryzalin (OZ) or 1µM Latrunculin B (LB) from simulations (above) and experimental data (below). **(C)** Kinetics of TGR of RG phase RHs in mock conditions (black line) or treatments with (orange line) either 10µM Taxol (TX) or 5µM Jasplakinolide (JP). **(D)** Kinetics of Nuclear distance from the tip of RG phase RHs in mock conditions (black line) or treatments (orange line) with either 10µM Taxol (TX) or 5µM Jasplakinolide (JP).

Simulation and experiments led to nucleus movement stabilisation after AF destabilisation-induced tip growth arrest, aligning with previous studies (*21*). Interestingly, this simulation of AF destabilisation in the G phase predicted a stagnation of the nucleus movement relative to the tip even after recovery, mirroring our experimental observations as a mean even though a high variability between samples was observed (fig. S7C). Either this stagnation of the nucleus results from a halt of the system after AF complete destabilisation, or it could be linked to an absence of vacuole movement resulting from its fragmentation (fig. S7D, E). The mean nuclear position remained further stable on average even after vacuole reformation, especially in G phase (fig. S7F), but looking at individual RHs showed chaotic nuclear movements upon vacuole reformation (fig. S7C), which is a good hint towards their interconnection, as shown by the large LR boxplot size (fig. S7F). This suggests a possible link between nucleus and vacuole movement during tip growth. Conversely, MT stabilisation by TX also caused tip growth arrest, characterized by the accumulation of AF bundles at the tip which was not reversed after drug removal (Fig. 4C, fig. S6D, see white arrow). This confirms that MT stabilisation in the RG phase is required for the RG/EM transition, whereas AF stabilisation had no significant effect on TGR during this phase (Fig. 4C, fig. S6E).

Overall, our data highlight the critical role of a timely MT and AF stabilisation in the subapical region for tip growth arrest during the RG/EM transition.

### Nuclear aspect ratio is regulated by both cytoskeleton and nucleoskeleton

Our simulations showed the nuclear AR to progressively increase from RG to EM, aligning with experiments (Fig. 2). To understand how this nuclear AR is regulated, we induced a destabilisation of either MT or AF in RG phase RHs, followed by a drug removal recovery phase, as previously described. Simulations predicted AR changes only upon AF destabilization, conflicting with experiments where nuclear AR was lower than in the control after both OZ and LB, even in recovery (Fig. 5A, B, n=6, p_OZ_ and p_LB_=0.02). This discrepancy may stem from balancing the behaviour with a G-RG-EM increase in AR in the model (Fig. 2G). Both AF stabilisation by JP (p=0.02) and MT stabilisation by TX (p=0.04) also led to a prevention of nuclear AR increase in late recovery, probably due to an increase of AF bundling around the nucleus and decrease of MT signal (fig. S6D, E). It thus appears that keeping both AF and MT at an average stability is necessary for the nuclear stretching process to occur during the RG phase. This further shows that MT and AF dynamics are mandatory for this nuclear elongation process. Indeed, in *fra2* mutants, constitutive MT stabilisation correlated with significantly lower nuclear AR than WT during RH development (fig. S8B). However, we cannot exclude other factors and counteracting effects, such as nucleoskeleton forces, which may contribute to cytoskeleton effects on nuclear AR.

**Fig. 5:**
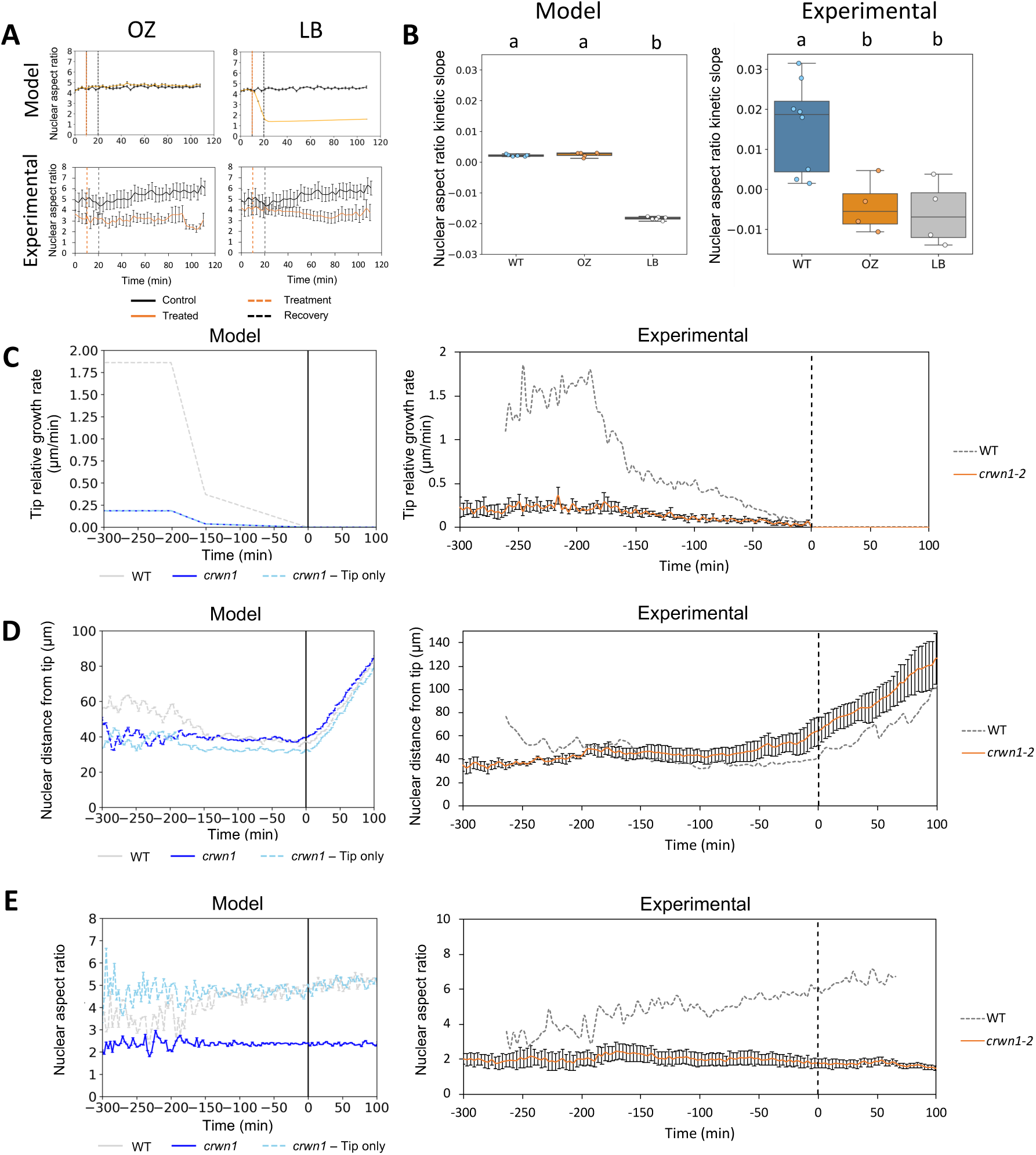
Behaviour of TGR and nuclear dynamics upon cytoskeleton drugs in RG phase RHs or mutation of the nucleoskeleton. **(A)** Mean nuclear aspect ratio of RHs in RG phase treated with either 1µM Oryzalin (OZ) or 1µM Latrunculin B (LB) from model (above) and experimental data (below). The kinetic of treated RHs (orange line) is composed of 10 min control medium, followed by 10 min of treatment, and 90 min of drug removal. The treatment is compared to the average TGR for control RHs (black line and left graph). (**B**) Mean nuclear velocity of RG phase RHs treated with OZ or LB, from model (left) or experimental data (right). Different letters highlight statistically different conditions according to Mann-Whitney non parametric test (p < 0.05). (**C-E**) TGR (C), nuclear distance from the tip (D), and nuclear AR (E) kinetic for *crwn1-2* mutant (orange line) in comparison with WT (dashed line) for both simulations (left) and experimental data (right). In simulation, we considered two scenarios: either i) additional nuclear spring force changes due to chromatin compaction and decreased nuclear area (blue line) or ii) reduced tip extensibility only (dashed blue line).

To test the effect of a constitutive impaired nuclear elongation in RH growth, we analysed the *crwn1-2* nucleoskeleton mutant of *CROWDED NUCLEI 1*, expressing SUN1-GFP. This mutant displays an impaired nuclear elongation (*47*) and shorter RHs compared to WT (fig. S8C) as well as an increase of AF bundling (fig. S8D). We replicated the observed *crwn1-2* reduced TGR (Fig. 5C, 0.4±0.1 µm/min, n=5) in our simulations (Fig. 5C) by decreasing tip extensibility but resembling the WT phase changes (Fig. 1). Assuming a timer-based cell size regulation (*48*), the reduction of tip extensibility led to shorter RHs in simulations and aligning with experimental data. Both simulations and experiments showed no difference in nuclei position to the tip in *crwn1-2* compared to WT in RG phase (Fig. 5D, n=5). In order to understand the lack of elongation of the nucleus in *crwn1-2,* we modelled two scenarios: i) reduced tip extensibility alone or ii) additional nuclear spring force changes due to chromatin compaction and a decreased nuclear area as previously reported (*43*) (text S1). Our predictions suggested that the constitutively lower nuclear AR in *crwn1-2* was primarily driven by the reduced nuclear area and stronger nuclear envelope spring forces (Fig. 5E), which we suggest may be due to an increase in AF bundling (fig. S8D). Despite a similar nuclear backward movement during EM in both *crwn1-2* and WT, the nuclear AR remained unchanged in *crwn1* (Fig. 5E, fig. S8E, G). Even though some faint stretching of the nucleus occurred during the G phase, the nucleus became more rounded as the tip growth arrested, persisting during nuclear repositioning (movie S7). This may be due to the loss of EMTs during RH maturation disrupting the nuclear-tip connection, as well as the inability of the nucleoskeleton to maintain the nuclear elongation initially driven by cytoskeletal forces. Indeed in *fra2*, the nuclear squeezing by MTs was alleviated in the EM phase, with both nuclear AR and area increasing (fig. S8B, F, G, H), resuming G phase levels from WT lines.

Overall, our findings suggest a cytoskeleton-nucleoskeleton interplay at the nuclear envelope to regulate the nuclear AR, which should be refined in our model, even though our *crwn1* model tends to show that these changes could be assured by nuclear mechanics alone. It further shows the link between nuclear compaction and tip growth rate.

### Vacuole movement and cell stiffness changes can provide causality to transitions between root hair growth phases

Our model could not fully reproduce the effects of AF destabilisation and recovery on nuclear distance from the tip in two main aspects : i) a high variability in the nuclear trajectory during the treatment recovery between G phase RHs and ii) a global backward nuclear movement in RG phase RHs. These results suggest that another organelle, such a the vacuole which is fragmented upon LB treatment, may influence nuclear movement during RH growth. To test this, we analysed vacuole positioning kinetics relative to the tip during RH growth (Fig. 6A, fig. S9A). During the G phase, the vacuole distance from the tip remained stable (10.1±0.2 µm) but significantly decreased in the RG phase (5.1±0.1 µm) (Fig. 6A, n_G_=46, n_RG_=36, p=7.9×10⁻¹⁵), eventually reaching the tip after tip growth arrest, as previously described (*49*). A statistical shift in vacuole distance was also observed upon G/RG (from 8.1±0.3 to 5.4±0.2 µm, p=0.01, Fig. 6B, fig. S9A). Inducing a G-to-RG transition-like with 1 µM OZ produced a similar vacuole shift, which persisted after drug recovery (Fig. 6C, n=6, p_C/T_=0.002; p_C/LR_=0.008). A comparable vacuole positioning pattern was observed in *crwn1-2* mutant RHs maintained in a RG phase (Fig 6D, 6.3±0.6 µm, n=10). Interestingly, in *fra2,* the nucleus shifted to RG values at tip growth arrest while vacuole position shifted to EM values, suggesting that the G/RG transition is still occurring in this mutant independently of tip growth dynamics.

**Fig 6:**
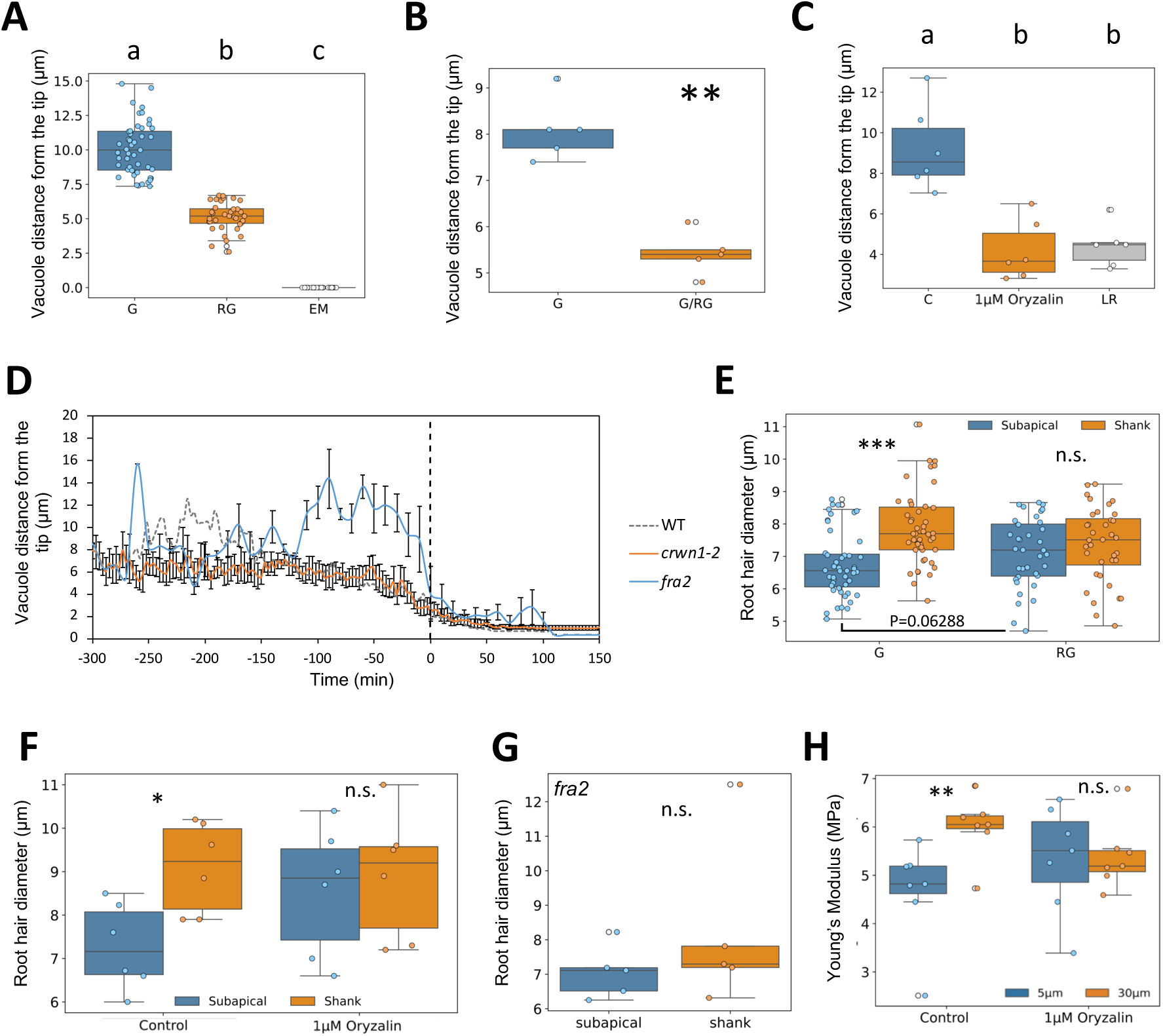
Changes in cell diameter, vacuole position and cell stiffness in the subapical region during RH growth. (**A-B**) Distribution of vacuole position from the tip at each developmental stages (**A**) and within the 20 min around the G to RG transition in WT (**B**). (**C**) Vacuole position from the tip upon 1µM Oryzalin treatment (T) and further late recovery from drug removal (LR) compared to G phase control (C). Different letters highlight statistically different conditions according to Mann-Whitney non parametric test (p < 0.05). (**D**) Kinetic graph of vacuole position compared to the tip in WT (black dashed line), *fra2* mutant (blue line) and *crwn1-2* (orange line). (**E-G**) RH diameters in the sub-apical and shank regions at fixed position from the tip, between G and RG phases (**E**), upon OZ treatment (**F**) and in *fra2* (**G**). (**H**) Cell stiffness in the sub-apical and shank regions upon OZ treatment compared to control. **: P-value <0.01; ***: P-value < 0.001.

Since the vacuole stabilizes at ∼5 µm from the tip during RG (proximal subapical region, hereafter called “subapical”), we examined RH diameter changes at this position during maturation, defining it as the “subapical” region. For comparison, we analysed a point at 30 µm from the tip in the proximal shank (hereafter called “shank”), where diameter changes were unlikely due to the presence of both primary and secondary cell walls. Using differential interference contrast (DIC) microscopy, we tracked diameter variations in these two regions. In the G phase, subapical RH diameter was smaller than in the shank (Fig. 6E; n_G_=46, d subapical=6.7±0.1 µm, d shank=7.9±0.2 µm, p=3.9 × 10⁻⁶) but they both gradually equalized with each other within -100 to -50 min before tip growth arrest (Fig. 6E, fig. S9B; n_RG_=36, p=0.3) due to a tendency for the subapical diameter to increase (p =0.06288). A similar equilibrium occurred after MT destabilisation with 1 µM Oryzalin during the G phase (Fig. 6F, n=6, p=0.03). In *fra2*, shank and subapical diameters showed no significant difference during G phase (Fig. 6G), possibly due to a defect in wall thickness as previously described (*45*) which could result in the observed low diameter in the shank. Given its connection to hydraulic status, vacuole positioning could also locally affect RH shape and cell wall stiffness.

To explore the link between diameter changes and cell stiffness, we used atomic force microscopy (AFM) coupled with fluorescence imaging (*50*). Using a large size tip, our AFM measurements are expected to reveal a combination of both cell wall elasticity and turgor pressure (*51*). The apparent Young’s modulus measurements in G-phase RHs revealed significant stiffness differences correlating with diameter variations (Fig. 6H, n=7, p=0.01). However, MT destabilisation with OZ cancelled this stiffness difference between the two regions (p=0.79), suggesting a relationship between MTs, RH diameter, and cell stiffness. Notably, we only observed an OZ-induced decrease of shank stiffness when the cell wall had been enzymatically digested for 5 min (fig. S9C, n=6, p=0.02). This suggests that the equilibration of RH diameter between shank and sub-apical regions mainly results from an adjustment of local cell stiffness between both regions, possibly driven by vacuole repositioning at G/RG. Cell stiffness in the shank was also drastically reduced in MRHs compared to GRHs (fig. S9D, n=5 p=0.04). These findings suggest that MTs regulate cell stiffness during RH growth, and the reduced CMT density in mature RHs may also contribute to the observed decrease in cell rigidity. Thus, a transient MT destabilisation triggers organelle reorganisation at G/RG, with the nucleus and vacuole shifting toward the tip, then remaining at a stable position, resulting in a local adjustment of cell stiffness.

## Discussion

Using the new imaging resolution brought by the microfluidic technology, we identified that tip growth in RHs goes through timely coordinated transitions between phases—namely, G to RG and RG to EM, the latter corresponding to tip growth arrest. Thus we can complete the RH development cycle which was previously described for early initiation (*52*). Our data implies transient changes in cytoskeletal stability and organisation which leads to changes in TGR dynamics during these developmental phases, in correlation with fast modifications of nuclear positioning and slow changes in its morphodynamics (Fig. 7). However, some discrepancies between our model and experiments may suggest that vacuole movement and cell mechanics are important for such transitions in addition to cytoskeleton reorganization.

**Fig. 7:**
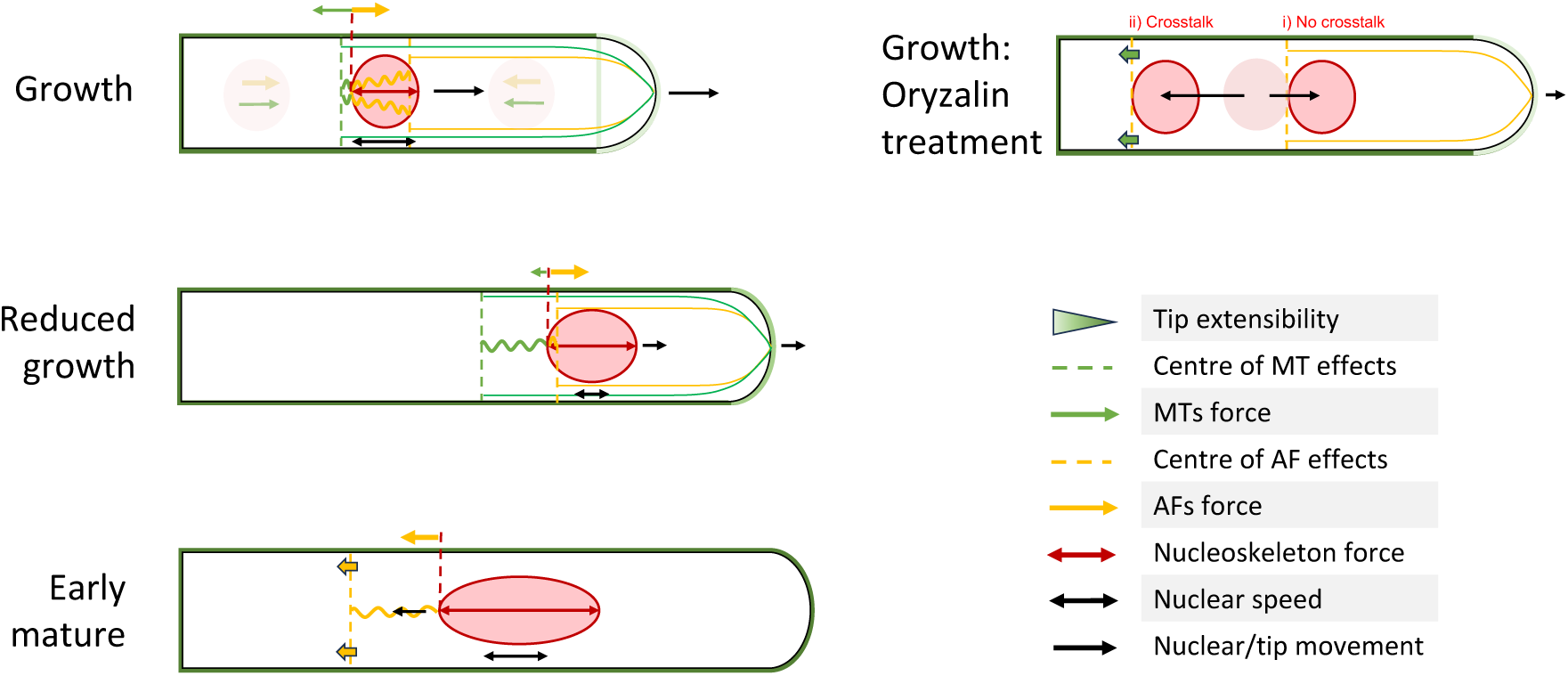
Integrative Model of RH development and growth phases. In Growth Phase: The tip advances at a speed of around 1-1.5µm/min, followed by the nucleus at around 60-70µm from the tip. Our model hypothesizes that AFs forces majorly maintain the connection of the nucleus to the tip (left side). However, if the nucleus position was further away from the centres of cytoskeleton forces it would move back towards them as to an equilibrium position (nuclei and forces in light colours). These AFs forces between the back of the nucleus and the centre of AFs effects at the front create compression forces resulting in nucleus squeezing, thus reducing nuclear apparent area. **In Growth phase with Oryzalin Treatment:** Two scenarios with (i) and without (ii) crosstalk of MTs on AFs are presented. i) With no crosstalk, after Oryzalin treatment in G phase the MT spring forces reduce to zero and the nucleus moves forwards and settles to the equilibrium position of the actin force slightly closer to the tip (pale red circle shows the nucleus old position and dark red circle its new position). ii) With crosstalk, after Oryzalin treatment, MTs crosstalk on AFs additionally causes the AFs connection to the tip to be weakened and the centre of the AFs force to retreat from the tip, pulling the nucleus away from the tip, resembling an EM phase. **In Reduced growth phase**: MT stability transiently decreases resulting in the MT fringe disappearance and in an increase of AF stabilization in the subapical region. This triggers a fast decrease in TGR, but G-actin at the tip maintains a state of growth (matching our simulation of reduced tip extensibility). The nucleus moves forward as a result of increased AF forces, stabilizing in front the of centre of these AF forces. The increase of AF stability also leads to a decrease in nuclear back and forth motion. Finally, our model displays a progressive increase of the nucleus AR from nucleus internal forces counterbalancing the reduction in AF compression forces. Coincidently, the AF cables progressively become more present around the nucleus through the whole stage. **In Early mature phase**: After a progressive stabilization of AFs in the apical region, the long-convoluted AF cables trigger tip growth arrest (matching our simulation of reduced tip extensibility to the maximum). Following this, the nucleus reaches its maximal elongation, supported by inner nucleoskeleton forces. The centre of AF forces moves backwards, resulting in a backward movement of the nucleus towards the RH base.

### Transitions between growth phases, a matter of timely modifications of cytoskeleton stability

We demonstrate that RH development deviates from the expected pattern, i.e. a high initiation growth rate quickly followed by a gradual deceleration until tip growth arrest(*48*). Instead, it involves a phase of rapid and constant elongation, followed by a shift in subcellular dynamics that induces a swift transition to a slower growth phase, during which the growth rate gradually declines in a timely fashion until reaching zero at tip growth arrest. This precise temporal regulation orchestrates the coordination of tip growth phases until growth arrest. Previous studies have shown that TGR decreases when RHs are grown in media of increased substrate stiffness, such as agar (*48*). On average, it is already below the 0.6µm/min threshold for agar concentrations above 0.5%, which could explain why the G/RG transition has not been observed in these conditions. Similarly, the presence of such a transition is not observed in the *crwn1-2* mutant where the TGR is already below this threshold at around 0.25 µm/min. Therefore, our microfluidic experimental setup appears to be a remarkable tool for capturing RH growth kinetics without any mechanical interference of the substrate, thus allowing the observation of this very subtle growth transition mechanism.

Different mechanisms for cell fate determination have been identified including size control theoretical strategies named sizer, timer and adder (*53*). For example, a cell size-mediated transition has been shown to be a robust mechanism for stomatal fate commitment (*54*). However, RH growth phase transitions appear to rely on a time-based regulatory mechanism instead. This is supported by our findings that G/RG always seem to occur at approximately 150 min before tip growth arrest, while the RG/EM transition in terms of cytoskeleton reorganisation and nucleus positioning appears to occur precisely at tip growth arrest. Further evidence for this time-based control comes from observations that RH growth time courses remain consistent across different agar concentrations with only the TGR maximum being affected (*48*). External clues can also modulate RH growth through the synthesis of the transcription factor RSL4 during the initiation of hair elongation. Such a factor is further gradually degraded during RH growth and can thus be used as a clock to limit RH development time (*52*).

### Tight coordination between tip growth, nuclear and cytoskeleton dynamics

During G/RG, we observed a decrease in MT intensities, along with the stabilisation and bundling of AFs in the subapical region. This coincided with the disappearance of the subapical MT fringe, which may be used as a scaffold for subapical/apical AFs, as shown in *Hydrocharis* RHs (Tominaga et al., 1996). The AF bundling likely limits vesicle trafficking for cell elongation(*44, 55*), thereby reducing TGR. This is supported by observations in the *vln1* mutant, where preventing AF bundling correlated with the maintenance of a high TGR (*56*). Similarly, MT stabilisation observed in *fra2* maintains a high TGR together with nuclear positioning, while a reduction in nucleus-tip distance occurs at the end of RH growth when EMTs disappear and CMTs signal decrease.

Such reduction in nucleus tip distance was mathematically predicted late in *fra2*, while very early in *crwn1* caused by the differences in TGR. This suggests that in WT conditions, the transition from G to RG implies a timely coordinated regulation of the stability of MT and AF networks. Notably, EMTs may be at play in this connection of the nucleus to the tip contributing to the coordination of TGR with nuclear positioning as reported in Medicago (*57*). Indeed, MT depolymerisation or stabilisation in Medicago resulted in a twofold reduction in TGR after 30 to 60 min of observation, resembling our results in *Arabidopsis*. This aligns with our mathematical model where a reduction in TGR was sufficient to cause a repositioning of the nucleus closer to the tip, due to the cytoskeletal forces linking nucleus and tip. Previous results in *Arabidopsis* more particularly had shown no effect of a depolymerisation of MTs after 10 min of treatment, which aligns with our results, as we show a significant effect of this treatment only in early recovery in the following 10 min. Thus, MT involvement for the coordination between TGR and nuclear positioning appears to be conserved across species. Additionally, this connection may rely on the involvement of MT motor proteins such as the kinesin-like ARK1 which is also involved in the maintenance of TGR, AF regulators localisation and AF-linked secretary molecules anterograde motility. This therefore supports the hypothesis of a MT influence on TGR via a crosstalk between MT and AF(*58*), which has only been linked to Myosin VIII so far in plant tip growing cells (*59*).

Our findings confirm that cytoskeleton-destabilizing treatments uncouple nuclear movement from TGR, highlighting the complexity of regulating cytoskeletal dynamics in RH growth phases. Indeed, we observed that the destabilisation of one cytoskeletal network triggers the reorganisation of the other, either during treatment or recovery phases, especially in the subapical region, highlighting the presence of an AF/MT cross-talk previously underestimated during RH growth. Regulatory proteins which bind both MTs and AFs, such as formins (*60*), whose activity of chemical inhibition has been shown to prevent nuclear migration in RHs (*61*), would be good targets to explore further for the understanding of such an interaction between both networks. In animals, the formin regulatory protein FHDC1 was shown to regulate both MT depolymerisation and AF polymerization, highlighting this potential dual role (*62*). In *Arabidopsis,* the overexpression of the formin FH8 was shown to result in shorter RHs due to a decreased AF severing activity (*63*). Finally, due to the presence of perinuclear MTs and an actin-myosin link to the nuclear envelope, cytoskeleton-induced changes in nuclear AR could also influence cytoskeletal stability and organization in the apical/subapical region for modulating TGR. In this context, the progressive increase in nuclear AR during the RG phase could lead to nuclear pore complex stretching (*64*), facilitating the nucleocytoplasmic shuttling of cytoskeletal regulators into the cytoplasm.

In this study, we provide evidence for the involvement of MTs extending from the nucleus to ensure its connection to the tip, probably through crosstalk with AFs (Fig. 7). The control of the cytoskeleton dynamics at the G/RG and RG/EM transitions may be regulated by the timely expression of cytoskeletal regulators. This, however, requires further investigation.

### Integration of cytoskeleton forces into the model

Our mathematical model successfully replicates many experimentally observed tendencies, including the maturation process, cytoskeletal depolymerisation treatments, and the effect of mutations. However, it shows these trends as much more significant than in our experimental data, highlighting the high variability between biological samples. This is not reproduced in simulations, where stochastic effects are restricted to a single noise term. Even so, by capturing these dynamics this model provides valuable insights into the complex interactions between cytoskeletal organization, RH growth, and nuclear movement and morphodynamics. However, we do not consider cytoskeleton stabilisation treatments at this stage, as a more detailed cytoskeletal study is required first to understand their effects on the fibre patterning and network forces in the RH, particularly in relation to tubulin availability and vesicle trafficking for tip growth.

A key uncertainty is the net direction of AF and MT forces on the nucleus, which for this model we assumed correlated to peaks in AF and MT fluorescence intensity relative to the nucleus position. During the RG phase we observed a reorganisation of the cytoskeleton and a reduction in MT density correlating with a forward movement of the nucleus. However, in Oryzalin treatment we observed a significant backward movement of the nucleus within the first 10 min following MTs depolymerisation. These differences are thus difficult to reconcile within our model assumptions but may indicate nuanced and intricate interactions between MTs and AFs. We are able to capture these contradictory nuclei movements through allowing for a cross-talk between AFs and MTs. This leads to a weakening of the AF connection to the tip upon MT depolymerisation which weakens the nucleus connection to the tip, thus resulting in the observed backward movement of the nucleus, although with a 20 min delay compared to experimental observations. Furthermore, in experiments the nucleus stabilizes at a new position upon MT repolymerization. This implies that AF forces may be actually directed toward both the RH base and the tip, with MTs primarily ensuring the nuclear connection to the tip through crosstalk with AFs rather than exerting a direct force on the nucleus. This aligns with previous studies (*18, 21*), indicating that MTs maintain the nuclear connection to the tip even in the absence of AF-myosin XI-i interactions. However, in our model the AF forces push the nucleus forward towards the tip, reflecting other studies that AF forces are the primary driver of nuclear movement in RHs (*19*). Comparison with these previous studies actually suggests a difference between the effect of fine AFs in the subapical region and AFs bundles in the shank. A destabilisation of fine AFs (but not of AF bundles) by Cytochalasin D results in a backward movement of the nucleus similar to a mature RH, suggesting a force of AF bundles directed backwards. Conversely, unbundling AFs using an anti-villin antibody—without causing depolymerization—results in a nuclear forward movement probably related to fine AFs forces at the tip. Furthermore, oryzalin-induced MT depolymerization, which enhances AF bundling, likely amplifies this backward force. This may explain the delayed nuclear backward movement which we observed with oryzalin treatment compared to the faster response seen with Cytochalasin D in the study by Ketelaar et al. (*19*). The fact that the nucleus is significantly displaced only after complete MT depolymerisation further underscores that changes in the balance of forces within the cellular environment are not immediate.

A balance between AF and MT forces along with nucleoskeleton forces is also necessary to account for changes in nuclear AR. In WT maturation we see a progressive increase of this AR through RG and EM phases which is reflected in the model as a consequence of the nucleus being released from AF squeezing effects while moving towards the tip. MT forces pull the nucleus in the opposite direction but are weaker than AF forces. Interestingly, the *crwn1-2* mutation also results in a decrease in nuclear AR compared to WT. In the model, this low AR is a result of reduced nuclear spring forces representing nucleoskeleton forces as well as nucleus volume, agreeing with experimental observations showing that this low AR also correlates with a lower nuclear area in *crwn1-2* mutant. One suggestion to explain this would be a mechanical feedback response mediated by LINC complexes and myosin XI-i (*65*). This phenotype might equally be due to an increase in heterochromatin compaction at the nuclear periphery, which has already been demonstrated in this mutant (*66*). Similar to *crwn1-2,* we observe a decreased nucleus area in *fra2* mutants compared to WT. However it increases back to a similar level as WT around maturation before dropping back again. This correlates with the decreasing MT signal observed in the late stage of growth in this mutant, similar to WT. MT over-stabilisation in *fra2* may initially compress the nucleus during early growth, reducing its apparent area. As MT nucleation may gradually decrease in later stages, this squeezing effect diminishes, allowing the nucleus to recover its resting size. This highlights a complex and intricate network controlling nucleus shape which still requires thorough study.

### Importance of cell mechanics in the control of tip growth

Beyond the cytoskeleton and nucleus, we also observed correlations between vacuole position, RH diameter, and cell stiffness as TGR decreases during the RG phase, as well as during the recovery phase following MT depolymerisation with Oryzalin. An increase in RH diameter and cell wall thickness has been associated with reduced tip growth, as seen in the *rdh4* mutant (*49*). This mirrors our observations in the RG phase within the proximal subapical region, suggesting a relationship between RH diameter and cell stiffness. Interestingly, the *eru* mutant, which exhibits a twofold increase in cell wall thickness, also displays a disrupted RH elongation (*67*). However, despite its thicker cell wall, microplate bending experiments revealed that cell stiffness in this mutant is lower than in WT (*22*), suggesting that the vacuole-linked turgor pressure likely plays a significant role. Our AFM data showed an apparent elastic modulus of approximately 5-6 MPa, consistent with previous studies using similar measurement conditions (*68*). After enzymatic digestion of the cell wall, cell stiffness values decreased 10-fold compared to the control. Such values aligned with previously reported turgor pressure (0.7 MPa in average) (*69*). Notably, a 10 min MT destabilisation caused a further decrease in cell stiffness, as shown previously on protoplast cells (*70*). Therefore, our results highlight a connection between MTs and RH cell stiffness, potentially mediated by osmotic changes which were shown to impact it (*71*). Another possible contributor to TGR reduction is that vacuole expansion in the proximal subapical region might hinder vesicle trafficking at the tip, thereby reducing exocytosis dynamics required for tip growth, as previously suggested (*40*). However, this remains to be experimentally validated. This vacuole expansion during the RG phase may also contribute to the stabilisation of the new nucleus position relative to the tip. During RH maturation, MT density decreases, coinciding with vacuole expansion into the apical region potentially correlated with the secondary cell wall reaching the tip (*49*). Since GRH cell stiffness is significantly higher than that of MRH, it suggests that turgor pressure may decrease upon RH maturation. Alternatively, we cannot exclude changes in cell wall thickness as secondary wall deposition was shown to cover the whole RH cell after maturation (*49*). To further explore these mechanisms, future work should focus on developing tools to track turgor pressure and cell wall stiffness throughout RH development in conjunction with live-cell imaging. Notably, plant zygote cells, which also exhibit apical growth, display growth kinetics which include changes in tip diameters and internal forces, suggesting a transient increase in turgor pressure modulates cellular growth stage and division (*4*). Modelling of this cell incorporating turgor pressure and cell wall extensibility, allowed a reconstruction of the changes observed experimentally, highlighting the timely coordination of subcellular elements for this particular type of apical growth, even though it differs significantly from what is observed in RHs (*72*).

In our mathematical model, wall mechanics were simplified for example by assuming a rigid cell wall outside the tip growth zone and neglecting vacuole effects. Despite its limitations this model captures many of the observed nucleus behaviours during the G to EM phases. However, some discrepancies were observed when comparing it to the experimental data, particularly in nucleus positioning relative to the tip under certain drug treatments. These experimental findings suggest different AF and MT force directions compared to our model and what has been previously proposed. We suggest that the interplay of these cytoskeleton forces with nuclear behaviour and morphodynamics is also related to vacuole dynamics during RH development. This relationship also appears to affect observed changes in cell stiffness and morphology. Future work could expand our mathematical model to incorporate vacuole effects then test its interplay with different AF and MT force directions. These could be more accurately tested using the technology of optical tweezers (*73*). However, to accurately incorporate vacuole effects a more detailed understanding of how it interacts with the different cellular components would first be necessary, potentially using a 3D finite element model.

## Material and Methods

### Plant material and microfluidic device setup

We have used *Arabidopsis thaliana* Col-0 seedlings expressing different fluorescent tag constructions. A line expressing *proSUN2:SUN2-tagRFP* (*65*) was crossed with another line expressing *pro35S:GFP-MBD* as described previously (*23*) to visualize both the nuclear envelope and the microtubule network. The same *proSUN2:SUN2-tagRFP* line was crossed in parallel with a line expressing *proUBI:LifeAct-GFP* (*74*) to visualize both the nuclear envelope and the AF network. The SUN marker line was introgressed into both mutant backgrounds *crwn1-2* (SALK_041774, (*47*) and *fra2* (EMS mutant), (*45*).

Design and assembly of the microfluidic chips are described in (*42*). Resulting seeds were sterilized and transferred into cut micropipette tips filled with ½ MS media (Duchefa Biochemie; Haarlem, The Netherlands) supplemented with 1% agar and 0.5% sucrose. The resulting tip assembly was stratified for 48hrs in the dark at 4°C and germinated under a 16h of light/8h of dark regime with a light intensity of 70 µmol.s^-1^.m^−2^ of fluorescent lighting at 23°C. After 5 days, the tip grown seedlings were transferred into a microfluidic chip under a constant flow rate (8 µl.min^−1^) of ½ MS liquid media supplemented with 0.5% sucrose as described (*23*) using a peristaltic Fusion 200 pump (Chemyx Inc., Stafford, TX, USA) and the seedlings were grown under the same light and temperature conditions as above. Live microscopic imaging of plant RHs grown in the micro-channels of the microfluidic chip was performed after 4 to 5 days of seedling growth (i.e. 9 – 10-day old seedlings).

### Drug treatment and confocal image acquisition and analysis

To study the effects of MT and AF networks on RH tip growth and nuclear position and shape in real-time, we supplemented the ½ MS liquid media circulating in the microfluidic chip with 1µM of Oryzalin (MT polymerisation inhibitor), 1µM of Latrunculin B (AF polymerisation inhibitor), 10µM of Taxol (MT stabilisation) or 5µM of Jasplakinolide (AF stabilisation). Liquid media supplemented with DMSO in proportions matching the quantities added with each drug was used for both control and recovery imaging. The change of media between control, treatment and recovery experiments was performed by changing the media-filled syringe on the pump, increasing the flow rate up to 1000 µl.min^−1^ for 45-60 seconds while starting the image acquisition, and then quickly reducing the flow rate back to 8 µl.min^−1^. The increase in flow rate was important to quickly replace the media in the microfluidic chip and avoid air bubble accumulation in the micro-channels upon the change of media.

Image acquisitions were performed with a LSM700 confocal microscope (Zeiss, Wetzlar, Germany) using an immersion oil 40×0.3 lens objective in multitracking mode. GFP was imaged with a 488 nm excitation and a 510 nm emission wavelength and tagRFP was imaged with a 555 nm excitation and 617 nm emission wavelength. Real-time image acquisitions were performed using a 1.0 µm resolution Z-stack every 3 min for 10 min for control and treatment, and for 90 min for the drug removal.

### 3D segmentation and intensity profiles

The 3D segmentations were performed using cv2 in Python on z-stack experimental images with a separate channel for the nucleus and either AFs or MTs. The nucleus is segmented using a two loop steepest ascent method with post processing. For the cytoskeleton, the signal is segmented in two ways and compared to identified sections of non-polymerized signal which are then removed. The 1D intensity profiles were obtained from the original images either by summing over the signal from all slices of the Z-stack or by z-projecting and taking the mean in each location, with the nucleus channel fluorescence intensity peaks used to determine the nucleus location. Manual pre-processing was necessary with some images to reduce background noise levels and fluorescence reflected from the microchannel.

### Image analysis

Image quantification and kinetics of nucleus and cytoskeleton behaviour was performed using ImageJ v.1.53q (NIH, Bethesda, MD, USA). TGR, nucleus and vacuole distance to the tip and RH diameter were manually measured using DIC and RFP channels. Nuclear shape was segmented from tagRFP Z-stacks, and nucleus morphological parameters were assessed from the resulting ROIs using ImageJ internal plugins. For dynamic profiles plots of cytoskeleton organization, RH endpoints were manually identified and annotated in each frame of the collected time series data for the alignment of all profiles in regards to the RH tip. Following the identification of these endpoints, longitudinal intensity profiles were extracted from RFP (nucleus) and GFP (AF and MT) fluorescent channels at each time point across the time series, using transversal averaging over RH width. These intensity profiles thus represent the variation in signal intensity along the longitudinal axis of the RHs.

### RH modelling

The RH surface is assumed to have a circular cross-section and to be rotationally symmetric about its long central axis. The tip region (a distance L from the apex) has a prescribed extensibility function allowing its growth, while the rest of the surface is stationary. Stress and strain are calculated from the pressurized cell shape, which then determines how the cell shape evolves over time of simulated growth (*27*). The nucleus is modelled as a spheroid with a fixed volume V and a resting aspect ratio that sets its shape in the absence of external forces. Two spring forces representing the AF and MTs act on the nucleus. An additional surface force representing the nucleoskeleton acts to restore the nucleus to its rest shape if it is perturbed. A normally distributed stochastic force also acts on the nucleus. Boundary forces restrict the nucleus to the cell interior and represent the nucleus physically pressing up against the cell membrane. As previous studies have indicated that AF forces are more important for nucleus movement in the growing RH (*21*), we set the parameter controlling AF effects to have a larger magnitude than the corresponding one for the MT effect. To simulate RH development we fit the tip extensibility to set the TGR and use changes in the AF and MT peak fluorescence intensities to set the AF and MT force parameters.

### Atomic Force Microscopy (AFM) indentation measurements

Indentation measurements were performed with a Bioscope II Catalyst Nanoscope V (Bruker, software version 9.0) coupled to a Leica DMI 6000 B inverted epi-fluorescence microscope. Seedlings of SUN2-tagRFP X Lifeact-GFP and SUN2-tagRFP X GFP-MBD lines grown for 7 days on ½ MS plates supplemented with 1% sucrose were mounted on a microscopic slide using a silicone Liveo^TM^ MG-2502 glue (Dupont, Mechelen, Belgium). The microscopic slide was then mounted with a drop of demineralized water on the microscope stage.

Measurements were performed under various conditions such as MT depolymersation by Oryzalin or a mild digestion of the cell wall. Oryzalin treatment was performed by replacing the drop of demineralized water on the slide by a solution of 1µM Oryzalin in demineralized water. For a mild digestion of the cell wall we used for 5 min a mix of enzymes [Cellulase 3% Onozuka R-10 (Merck, Saint-Quentin-Fallavier Cedex, France), macerozyme 1% (Duchefa Biochimie, Haarlem, The Netherlands), pectolyase 0.2% (Kyowa Chemical, Tokyo, Japan)], in MES 25 mM, CaCl2 8 mM, Mannitol 600 mM (Duchefa Biochimie, Haarlem, The Netherlands), pH 5.5 – diluted 1/10 in PBS. We then removed the digestion solution and replaced it by PBS prior to the indentation measurements.

The measurements were done with a high density carbon spherical tip 500 nm in radius, mounted on a cantilever with a 40 N/m spring constant (Biosphere, Nanotools). The force axis was calibrated by first measuring the deflection sensitivity of the photodiode. Briefly a force curve was acquired on a sapphire disk: on such material and considering the forces exerted, indentation is negligible, so the deflection sensitivity, linking the actual deflection of the cantilever to the output of the photodiode, is just the inverse of the slope of the curve after tip contact. Then the spring constant of the cantilever was measured by the standard thermal noise method. At each new experiment, the spring constant measured the first time was used, while the deflection sensitivity was measured each time.

The RH was scanned in PeakForce Tapping to acquire its topography. A series of force curves were performed along the main axis of the RH from 5 µm up to 35 µm from the RH tip. Force curves were performed with the following parameters: ramp size 2 µm, ramp speed 4 µm/s, setpoint force 1 µN. Quantification of the Young’s modulus for each curve was done by using processing software Nanoscope Analysis 2.0 (Bruker). Force vs. distance curves were first flattened by removing the result of a linear fit to the non-contact part of the force curve (Baseline correction), in order to set this part to 0 force. The force vs. tip-sample distance was then obtained calculating a new axis of distances as height Z – cantilever deflection Δd. Young’s modulus was obtained by fitting the extended segment of force vs. tip-sample distance curves with a Linearized Hertz model setting a tip radius of 500 nm and a Poisson ratio n of 0.5 (incompressible material). Resulting curves of force were processed and analyzed using the AFM image analysis software using the Hertz’s model to estimate the Young’s modulus for each indentation.

### Statistical analysis

The collected data were analysed for statistical significance using analysis of variance (one-way ANOVA with post hoc Tukey’s HSD test) and paired or unpaired *t*-test, two-tailed at *P* < 0.001, *P* < 0.01, and *P* < 0.05. Data were tested for statistical differences using a non-parametric Mann-Whitney U-test, with a p-value of 0.05. Modelling results were similarly tested for significance using the two sides Mann-Whitney U-test at 0.05 significance level, using the mannwhitneyu package in scipy.stats in Python3. All data are shown as the means with standard error (mean ± SD/√$).

## Acknowledgments

We thank F. Cvrckova for providing us the ubi-Lifeact-GFP line, K. Tamura for the SUN2-tagRFP line and France Bio imaging for microscopy facilities. We thank E. Hoffmann for technical assistance.

## Authors contributions

Conceptualization: M.E.C., H. J. and G.D; Data analysis :T. S., G. D., M. E. C., Funding acquisition: M.E.C., H. J., O. H. ; Methodology: G.D., G. S. , T. S., S. B., J. M., A. B. ; Investigation : G.D., G. S., S. B., T. S., S. B. ; Supervision: M.E.C., H. J.; Writing - original draft: G.D. , M.E.C., review and editing: G. D., M.E.C, H. J., T. S., O.H., E. H.

## Data and materials availability

All data needed to evaluate the conclusions in the paper are present in the paper and/or the Supplementary Materials. Data from the mathematical modelling will be deposited on our Gitlab webserver : https://gitlab.developers.cam.ac.uk/slcu/teamhj/publications/dupouy_spelman_etal_2025.

## Bibliography

1. J. L. Goldberg, How does an axon grow? Genes Dev 17, 941–958 (2003).

2. J. P. Bibeau, G. Galotto, M. Wu, E. Tuzel, L. Vidali, Quantitative cell biology of tip growth in moss. Plant Mol Biol 107, 227–244 (2021).

3. P. J. Brown, D. T. Kysela, Y. V. Brun, Polarity and the diversity of growth mechanisms in bacteria. Semin Cell Dev Biol 22, 790–798 (2011).

4. Z. Kang et al., Coordinate Normalization of Live-Cell Imaging Data Reveals Growth Dynamics of the Arabidopsis Zygote. Plant Cell Physiol 64, 1279–1288 (2023).

5. M. N. de Keijzer, A. M. C. Emons, B. M. Mulder, Modeling Tip Growth: Pushing Ahead. (Springer, Berlin, Heidelberg. , ed. Root Hairs. Plant Cell Monographs, 2009), vol. 12.

6. N. J. Brewin, Plant Cell Wall Remodelling in the Rhizobium–Legume Symbiosis. Critical Reviews in Plant Sciences 23, 293–316 (2004).

7. A. Mendrinna, S. Persson, Root hair growth: it’s a one way street. F1000Prime Rep 7, 23 (2015).

8. M. Akkerman et al., Texture of cellulose microfibrils of root hair cell walls of Arabidopsis thaliana, Medicago truncatula, and Vicia sativa. J Microsc 247, 60–67 (2012).

9. K. Herburger, S. Schoenaers, K. Vissenberg, J. Mravec, Shank-localized cell wall growth contributes to Arabidopsis root hair elongation. Nat Plants 8, 1222–1232 (2022).

10. T. Hirano et al., The SYP123-VAMP727 SNARE complex delivers secondary cell wall components for root hair shank hardening in Arabidopsis. Plant Cell 35, 4347–4365 (2023).

11. C. Ambrose, G. O. Wasteneys, Microtubule initiation from the nuclear surface controls cortical microtubule growth polarity and orientation in Arabidopsis thaliana. Plant Cell Physiol 55, 1636–1645 (2014).

12. N. Van Bruaene, G. Joss, P. Van Oostveldt, Reorganization and in vivo dynamics of microtubules during Arabidopsis root hair development. Plant Physiol 136, 3905–3919 (2004).

13. T. Ketelaar, N. C. de Ruijter, A. M. Emons, Unstable F-actin specifies the area and microtubule direction of cell expansion in Arabidopsis root hairs. Plant Cell 15, 285–292 (2003).

14. X. He, Y. M. Liu, W. Wang, Y. Li, Distribution of G-actin is related to root hair growth of wheat. Ann Bot 98, 49–55 (2006).

15. L. A. Vazquez et al., Actin polymerization drives polar growth in Arabidopsis root hair cells. Plant Signal Behav 9, e29401 (2014).

16. T. N. Bibikova, E. B. Blancaflor, S. Gilroy, Microtubules regulate tip growth and orientation in root hairs of Arabidopsis thaliana. Plant J 17, 657–665 (1999).

17. A. Paradez, A. Wright, D. W. Ehrhardt, Microtubule cortical array organization and plant cell morphogenesis. Curr Opin Plant Biol 9, 571–578 (2006).

18. C. Lloyd, K. J. Pearce, D. J. Rawlins, R. W. Ridge, P. J. Shaw, Endoplasmic microtubules connect the advancing nucleus to the tip of legume root hairs, but F-actin is involved in basipetal migration. Cytoskeleton 8, 27–36 (1987).

19. T. Ketelaar et al., Positioning of nuclei in Arabidopsis root hairs: an actin-regulated process of tip growth. Plant Cell 14, 2941–2955 (2002).

20. E. Chytilova et al., Nuclear dynamics in Arabidopsis thaliana. Mol Biol Cell 11, 2733–2741 (2000).

21. J. M. Brueggeman, I. A. Windham, A. Nebenfuhr, Nuclear movement in growing Arabidopsis root hairs involves both actin filaments and microtubules. J Exp Bot 73, 5388–5399 (2022).

22. D. Pereira, T. Alline, S. Schoenaers, A. Asnacios, In vivo measurement of the Young’s modulus of the cell wall of single root hairs. Cell Surf 9, 100104 (2023).

23. G. Singh et al., Real-time tracking of root hair nucleus morphodynamics using a microfluidic approach. Plant J 108, 303–313 (2021).

24. I. Meier, LINCing the eukaryotic tree of life - towards a broad evolutionary comparison of nucleocytoplasmic bridging complexes. J Cell Sci 129, 3523–3531 (2016).

25. K. Tamura, C. Goto, I. Hara-Nishimura, Recent advances in understanding plant nuclear envelope proteins involved in nuclear morphology. J Exp Bot 66, 1641–1647 (2015).

26. O. Campas, L. Mahadevan, Shape and dynamics of tip-growing cells. Curr Biol 19, 2102–2107 (2009).

27. J. Dumais, S. L. Shaw, C. R. Steele, S. R. Long, P. M. Ray, An anisotropic-viscoplastic model of plant cell morphogenesis by tip growth. Int J Dev Biol 50, 209–222 (2006).

28. E. Eggen, M. Niels de Keijzer, B. M. Mulder, Self-regulation in tip-growth: the role of cell wall ageing. J Theor Biol 283, 113–121 (2011).

29. M. Dogterom, S. Leibler, Physical aspects of the growth and regulation of microtubule structures. Phys Rev Lett 70, 1347–1350 (1993).

30. R. J. Hawkins, S. H. Tindemans, B. M. Mulder, Model for the orientational ordering of the plant microtubule cortical array. Phys Rev E Stat Nonlin Soft Matter Phys 82, 011911 (2010).

31. C. Gibson, H. Jönsson, T. A. Spelman, A mean-field theory approach to 3D nematic phase transitions in microtubules. https://arxiv.org/abs/2112.06855, (2023).

32. E. E. Deinum, S. H. Tindemans, J. J. Lindeboom, B. M. Mulder, How selective severing by katanin promotes order in the plant cortical microtubule array. Proc Natl Acad Sci U S A 114, 6942–6947 (2017).

33. F. J. Nedelec, T. Surrey, A. C. Maggs, S. Leibler, Self-organization of microtubules and motors. Nature 389, 305–308 (1997).

34. B. Rupp, F. Nedelec, Patterns of molecular motors that guide and sort filaments. Lab Chip 12, 4903–4910 (2012).

35. P. Krupinski et al., A Model Analysis of Mechanisms for Radial Microtubular Patterns at Root Hair Initiation Sites. Front Plant Sci 7, 1560 (2016).

36. M. Scianna, L. Preziosi, Modeling the influence of nucleus elasticity on cell invasion in fiber networks and microchannels. J Theor Biol 317, 394–406 (2013).

37. I. D. Estabrook, H. R. Thiam, M. Piel, R. J. Hawkins, Calculation of the force field required for nucleus deformation during cell migration through constrictions. PLOS Computational Biology 17, e1008592 (2021).

38. T. Hirano et al., PtdIns(3,5)P(2) mediates root hair shank hardening in Arabidopsis. Nat Plants 4, 888–897 (2018).

39. D. J. Cosgrove, Expansive growth of plant cell walls. Plant Physiol Biochem 38, 109–124 (2000).

40. M. E. Galway, J. W. Heckman, Jr., J. W. Schiefelbein, Growth and ultrastructure of Arabidopsis root hairs: the rhd3 mutation alters vacuole enlargement and tip growth. Planta 201, 209–218 (1997).

41. A. M. Emons, T. Ketelaar, in Root Hairs., A. M. Emons, T. Ketelaar, Eds. (Springer Berlin / Heidelberg, web, 2008), vol. 12, pp. 27–44.

42. G. Dupouy et al., Microfluidics to Follow Spatiotemporal Dynamics at the Nucleo-Cytoplasmic Interface During Plant Root Growth. Methods Mol Biol 2873, 223–245 (2025).

43. H. Wang, T. A. Dittmer, E. J. Richards, Arabidopsis CROWDED NUCLEI (CRWN) proteins are required for nuclear size control and heterochromatin organization. BMC Plant Biol 13, 200 (2013).

44. W. Pei, F. Du, Y. Zhang, T. He, H. Ren, Control of the actin cytoskeleton in root hair development. Plant Sci 187, 10–18 (2012).

45. D. H. Burk, B. Liu, R. Zhong, W. H. Morrison, Z. H. Ye, A katanin-like protein regulates normal cell wall biosynthesis and cell elongation. Plant Cell 13, 807–827 (2001).

46. M. R. Bubb, I. Spector, B. B. Beyer, K. M. Fosen, Effects of jasplakinolide on the kinetics of actin polymerization. An explanation for certain in vivo observations. J Biol Chem 275, 5163–5170 (2000).

47. C. Goto, K. Tamura, Y. Fukao, T. Shimada, I. Hara-Nishimura, The Novel Nuclear Envelope Protein KAKU4 Modulates Nuclear Morphology in Arabidopsis. Plant Cell 26, 2143–2155 (2014).

48. D. Pereira, T. Alline, L. Cascaro, E. Lin, A. Asnacios, Mechanical resistance of the environment affects root hair growth and nucleus dynamics. Sci Rep 14, 13788 (2024).

49. M. E. Galway, D. C. Lane, J. W. Schiefelbein, Defective control of growth rate and cell diameter in tip-growing root hairs of the rhd4 mutant of Arabidopsis thaliana. Canadian Journal of Botany 77, 494–507 (1999).

50. R. Goswami et al., Mechanical Shielding in Plant Nuclei. Curr Biol 30, 2013–2025 e2013 (2020).

51. S. Tsugawa et al., Elastic shell theory for plant cell wall stiffness reveals contributions of cell wall elasticity and turgor pressure in AFM measurement. Sci Rep 12, 13044 (2022).

52. S. Datta, H. Prescott, L. Dolan, Intensity of a pulse of RSL4 transcription factor synthesis determines Arabidopsis root hair cell size. Nat Plants 1, 15138 (2015).

53. N. Rhind, Cell-size control. Curr Biol 31, R1414–R1420 (2021).

54. Y. Gong et al., A cell size threshold triggers commitment to stomatal fate in Arabidopsis. Sci Adv 9, eadf3497 (2023).

55. D. D. Miller, N. C. A. De Ruijter, T. Bisseling, A. m. C. Emons, The role of actin in root hair morphogenesis: studies with lipochito-oligosaccharide as a growth stimulator and cytochalasin as an actin perturbing drug. The Plant Journal 17, 141–154 (1999).

56. N. Wang et al., The plant nuclear lamina disassembles to regulate genome folding in stress conditions. Nat Plants 9, 1081–1093 (2023).

57. B. J. Sieberer, A. C. Timmers, F. G. Lhuissier, A. M. Emons, Endoplasmic microtubules configure the subapical cytoplasm and are required for fast growth of Medicago truncatula root hairs. Plant Physiol 130, 977–988 (2002).

58. M. W. Yoshida, M. Hakozaki, G. Goshima, Armadillo repeat-containing kinesin represents the versatile plus-end-directed transporter in Physcomitrella. Nat Plants 9, 733–748 (2023).

59. S. Z. Wu, M. Bezanilla, Actin and microtubule cross talk mediates persistent polarized growth. J Cell Biol 217, 3531–3544 (2018).

60. F. Bartolini, G. G. Gundersen, Formins and microtubules. Biochim Biophys Acta 1803, 164–173 (2010).

61. S. Zhang et al., The migration direction of hair cell nuclei is closely related to the perinuclear actin filaments in Arabidopsis. Biochem Biophys Res Commun 519, 783–789 (2019).

62. C. S. Tong et al., Collective dynamics of actin and microtubule and its crosstalk mediated by FHDC1. Front Cell Dev Biol 11, 1261117 (2023).

63. K. Yi et al., Cloning and functional characterization of a formin-like protein (AtFH8) from Arabidopsis. Plant Physiol 138, 1071–1082 (2005).

64. C. E. Zimmerli et al., Nuclear pores dilate and constrict in cellulo. Science 374, eabd9776 (2021).

65. K. Tamura et al., Myosin XI-i links the nuclear membrane to the cytoskeleton to control nuclear movement and shape in Arabidopsis. Curr Biol 23, 1776–1781 (2013).

66. A. Poulet et al., The LINC complex contributes to heterochromatin organisation and transcriptional gene silencing in plants. J Cell Sci 130, 590–601 (2017).

67. S. Schoenaers et al., The Auxin-Regulated CrRLK1L Kinase ERULUS Controls Cell Wall Composition during Root Hair Tip Growth. Curr Biol 28, 722–732 e726 (2018).

68. M. Shibata et al., Trihelix transcription factors GTL1 and DF1 prevent aberrant root hair formation in an excess nutrient condition. New Phytol 235, 1426–1441 (2022).

69. C. Municio-Diaz et al., Mechanobiology of the cell wall - insights from tip-growing plant and fungal cells. J Cell Sci 135, (2022).

70. P. Durand-Smet et al., A comparative mechanical analysis of plant and animal cells reveals convergence across kingdoms. Biophys J 107, 2237–2244 (2014).

71. S. N. Shabala, R. R. Lew, Turgor regulation in osmotically stressed Arabidopsis epidermal root cells. Direct support for the role of inorganic ion uptake as revealed by concurrent flux and cell turgor measurements. Plant Physiol 129, 290–299 (2002).

72. Z. Kang, et al., A viscoelastic-plastic deformation model of hemisphere-like tip growth in Arabidopsis zygotes. Quant Plant Biol 5, e13 (2024).

73. M. Baclayon et al., Optical Tweezers-Based Measurements of Forces and Dynamics at Microtubule Ends. Methods Mol Biol 1486, 411–435 (2017).

74. F. Cvrckova, D. Oulehlova, A new kymogram-based method reveals unexpected effects of marker protein expression and spatial anisotropy of cytoskeletal dynamics in plant cell cortex. Plant Methods 13, 19 (2017).

