## Supplementary Figures, Text and Tables for "Root hair growth phases are coordinated by cytoskeleton, nucleus dynamics and cell mechanics in Arabidopsis"

### Supplementary Text S1: Details of the computational model

The tip-growing cell model refines the results from Dumais et. al [1] with stress-driving mechanical growth via biological characteristics to the *Arabidopsis* RH cell.

We assume the RH cell is symmetric around its long axis so it can be generated via a surface of revolution. Thus, our model only needs to track a single line on the cell surface coplanar to the centerline axis of revolution. For simplicity, we take the cell centerline as the x-axis in Cartesian coordinates, which can therefore define the cell surface line in the x-y plane where the y-coordinate also defines the local cross-sectional radius of the cell.

The cell surface (line) is discretized with points $(x_i, y_i)$ with the tip at $i = 0$ . The cell surface is split into two regions, a growth zone within distance $L$ of the tip along the cell surface consisting of $N(L)$ points and a non-growth zone further from the tip. $N$ is set to ensure numerical stability in the growth zone. As stress and strain are defined relative to the cell surface, we also use $s$ the distance along the demarcation line and $\theta$ the angle around the cross-section.

From the current cell surface shape, local curvatures $\kappa_s$ and $\kappa_\theta$ can be evaluated where subscript s denotes in the meridional direction and subscript $\theta$ denotes in the circumferential direction. From this, local cell surface stresses are given by

$$\sigma_s = \frac{P}{2\delta\, \kappa_\theta},$$

$$\sigma_\theta = \frac{P}{2\delta\, \kappa_\theta}\left(2 - \frac{\kappa_s}{\kappa_\theta}\right),$$

for cell properties pressure difference $P$ (between the inside and outside of the cell) and cell wall thickness $\delta$ [1].

We define cell wall extensibility $\Phi(s)$ as

$$\Phi(s) = \phi\, cos\left(\frac{s\pi}{2L}\right)^b,$$

taking $b = 10$ here and varying the extensibility magnitude $\phi$ over the simulation to account for different stages of RH growth. The extensibility decreases to zero away from the tip. This can then be used to define non-zero strain rates [1] as

$$\dot{\epsilon}_s = \Phi\,(\sigma_e - \sigma_y)\,\frac{(\sigma_s - \nu\,\sigma_\theta)}{K},$$

$$\dot{\epsilon}_\theta = \Phi\,(\sigma_e - \sigma_y)\,\frac{(\sigma_\theta - \nu\,\sigma_s)}{K},$$

when the effective stress $\sigma_e = [(\sigma_s - \sigma_\theta)^2 + (\sigma_\theta)^2 + (\sigma_s)^2]^{1/2}/\sqrt{2}$ is greater than or equal to the local yield stress $\sigma_y$ and where $\nu$ is a ratio determining how local strain is split in normal and tangent directions with

$$K = \left(\beta\,\sigma_s^2 + \beta\,\sigma_\theta^2 + (\beta - 6\nu)\sigma_s\sigma_\theta\right)^{\{\frac{1}{2}\}},$$

$$\beta = 2\,\nu^2 - 2\nu + 2.$$

The surface velocities in the normal $v_n$ and tangential $v_t$ direction can then be calculated as

$$\frac{\partial v_t}{\partial s} + \kappa_s\, v_n = \dot{\epsilon}_s\,,$$

$$\frac{cos(\Theta)}{r} v_t + \kappa_\theta\, v_n = \dot{\epsilon}_\theta\,,$$

for $\Theta$ the angle between the normal and cell surface and $r$ the cross-sectional radius. For a more detailed description see Dumais et. al. [1].

We use finite difference to discretise the cell surface (line) and solve the resulting system of linear equations for the velocity at every surface point. The surface can then be evolved forward a time $\delta t$ followed by a re-mesh of the surface if necessary.

The cell surface starts from an initial state (representing the trichoblast cell). We prescribe the initial cell surface as a sphere but this shape has minimal effect on our results provided it starts with a slight curvature to allow tip growth initiation.

**Nucleus Model**

The cell nucleus is modeled as a spheroid of constant volume $V$, which remains centered on the cell center line (i.e. the x-axis). We assume that forces are only applied at the front and rear of the spheroid (and as default apply them only at the back closest to the cell base). Thus, we only calculate the distance from the RH tip at time $t$ to the front (indicated by subscript $f$) and back (indicated by subscript $b$) of the nucleus. The distance to the front and back of the nucleus from the tip are $Z_f(t)$ and $Z_b(t)$, respectively (where $Z_b > Z_f$). The spheroid therefore has one semi-axis of length $(Z_b - Z_f)$ with the other two of length $R$ given by $R = \sqrt{3V/[4\pi(Z_b - Z_f)]}$. These three semi-axis lengths together determine the 3D spheroid nucleus shape. The distance of the tip from the RH base along its centerline is defined as $Y(t)$. Then $X_f(t) = Y(t) - Z_f(t), X_b(t) = Y(t) - Z_b(t)$ defines the exact position from the RH base of the front and back of the nucleus along the centerline.

The nucleus is modeled as being linked to the tip of the cell by two spring forces, one representing the actin forces of strength $A_a$ with equilibrium position a distance $B_a$ from the cell tip and the second representing the microtubule forces of strength $A_m$ with equilibrium position a distance $B_m$ from the tip. There is an additional internal spring force between the front and back of the nucleus of strength $\alpha_N$ preventing the nucleus from collapsing with rest length of $\beta_N$. This rest length (representing the preferred nucleus width) is split to have a cytoskeleton-dependent $\beta_{Nc}$ and cytoskeleton independent $\beta_{Nb}$ component so $\beta_N = \beta_{Nb} + \beta_{Nc}(A_a \,/\, \tilde{A}_a)$, where $\tilde{A}_a$ is a reference actin strength taken at cell initiation.

Incorporating these three forces, we take the movement of the front and rear of the nucleus as

$$\frac{\partial Z_f}{\partial t} = c - 2A_m\left(Z_f(t) + \tau_m(t) - B_m\right) - 2\,A_a\left(Z_f(t) + \tau_a(t) - B_a\right)$$
$$+ \left(X_b(t) - X_f(t) - \beta_N\right) + f(t, Z_f) + g(t, Z_f),$$
$$\frac{\partial Z_b}{\partial t} = c - 2A_m\left(Z_b(t) + \tau_m(t) - B_m\right) - 2\,A_a\left(Z_b(t) + \tau_a(t) - B_a\right)$$
$$-\alpha_N\left(X_b(t) - X_f(t) - \beta_N\right) + f(t, Z_b) + g(t, Z_b),$$

where $f$ is a normally distributed noise term; $g$ is a boundary force which is only non-zero when the nucleus is very close to the tip or base of the RH but keeps the nucleus within the cell; and $\tau_a(t), \tau_m(t)$ account for changes in the length of the connection between nucleus and tip. Note that the forces indicated by the second two terms are standard springs of length $B_a + \tau_a$ and $B_m + \tau_m$ respectively and if $\tau_a = \tau_m = 0$ the springs have a fixed length of $B_a$ and $B_m$ connecting the nucleus (equilibrium) position to the tip, i.e. the position traces the tip growth. The functions $\tau_a$ and $\tau_b$ account for cytoskeleton dependent changes in the connection between the nucleus and tip during cell growth, in physics terms for the spring rest length to increase, and introduced given the cytoskeletal data. These allow the locations of the cytoskeleton distributions (force potentials) to change leading to the nucleus to be pulled to a different location within the cell. They are modeled as

$$\frac{\partial \tau_a}{\partial t} = c\left(\frac{A_a}{\tilde{A}_a} - \frac{A_a}{\tilde{A}_a} A_{ct}\, e^{-C_{ct}\left(\frac{A_m}{\tilde{A}_m}\right)^{C_{ctf}}} - 1\right),$$

$$\frac{\partial \tau_m}{\partial t} = c\left(\frac{A_m}{\tilde{A}_m} - 1\right),$$

where $\tilde{A}_m$ and $\tilde{A}_a$ are reference actin and microtubule strengths taken as the values of $A_m$ and $A_a$ at tip growth initiation (corresponding to the start of the simulation). Note that when $\tilde{A}_m = A_m$ and $\tilde{A}_a = A_a$ then $\tau_a = \tau_m = 0$. This changes the spring forces relative to the tip when the actin or microtubule peak intensities change relative to their initiation value. The exponential term accounts for the cross-talk of microtubules on actin. A similar term could be added to the microtubule potential to account for actin on microtubule crosstalk but that is not considered in this paper. The exponential coefficients are taken large enough so that when $\tilde{A}_m = A_m$ and $\tilde{A}_a = A_a$, the exponential term is negligible.

The noise term $f$ is normally distributed with mean 0, with a standard deviation scaled relative to the microtubule and actin strengths. Specifically

$$f = N\left(0, \alpha^{\square}\left\{\beta + (1-\beta)\frac{A_a}{\tilde{A}_a}\left[\gamma + (1-\gamma)\frac{A_m}{\tilde{A}_m}\right]\right\}^2\right),$$

where $0 \leq \beta \leq 1$ defines the proportion of the variance dependent on actin strength, and $0 \leq \gamma \leq 1$ defines the proportion of the variance dependent on the microtubule strength, and $\alpha$ the initial magnitude of its standard deviation.

The boundary terms in $g$ are Heaviside functions which are non-zero when the nucleus approaches too close to the tip, base or sides of the RH. These represent the physical force of the cell wall on the nucleus when they are close enough to be directly interacting. These are defined as

$$g(t, Z_i) = \zeta\, H(\eta - X_i(t) - 1) tanh(\eta - X_i(t) - 1) - \zeta\, H(\eta - Z_i(t)) tanh(\eta - Z_i(t))$$
$$\pm \zeta_N\, H(\eta_N - |X_f(t) - X_b(t)|)\, tanh(\eta_N - |X_f(t) - X_b(t)|),$$

where $H$ is the Heaviside step function, $\zeta$ a constant determining the maximum magnitude of the boundary force at the RH tip and base, $\eta$ the interaction distance at the RH tip and base, $\eta_N$ the minimum nucleus length when the nucleus starts interacting with the cell side walls and $\zeta_N$ the maximum magnitude of this force. The subscript $i = \{b, f\}$ indicates the back or front of the nucleus respectively, with the positive final term taken for the back of the nucleus and minus for the front. The first term represents the RH base and nucleus interacting, the second term the nucleus and tip interacting and the third term the nucleus interacting with the sides of the cell. The tanh function ensures the force ramps up continuously when the nucleus is close enough to the boundary, while the Heaviside ensures 0 contribution outside of the boundary region. The additional $-1$ in the first term compared to the other is a numerical integration subtlety ensuring that we take the base cell of non-dimensional size of 1 so the back of the initiation cell is at $x = -1$. Note that the nucleus is rarely close enough to the tip or base of the RH for the first two terms to be non-zero, so their impact is believed to be minimal but they are included for completeness.

**Numerical solution of coupled nucleus-cell model**

We non-dimensionalize by the width of the starting cell (sphere) and solve the non-dimensional equations. The nucleus and tip are primarily linked by the tip position determined by the cell model which becomes the input for the nucleus model. They are solved in two steps. First, the cell model evolves the cell surface. The nucleus model works on a finer grid and calculates $c$ for the level of cell growth from $t_n$ to $t_{n-1}$ taking $Y_n(t) = Y_{n-1}(t) + ct$. The nucleus equations are then integrated. This process is then repeated.

The model was run in Python3 and the code is available open source from our Gitlab repository https://gitlab.developers.cam.ac.uk/slcu/teamhj/publications/dupouy_spelman_etal_2025.

Additionally, parameters are linked together to model particular behaviors such as cytoskeleton perturbations. Cytoskeleton forces are incorporated in the tip-growing cell model by associating particular mechanical parameters to cytoskeleton effects and in the nucleus model directly by linking those forces.
A list of parameters is shown in the Table S1. The parameters shown are non-dimensional. For our results, we redimensionalize with a non-dimensional distance unit representing 9 $\mu m$ and a non-dimensional time unit representing 0.5min. This scales the initial width of the cell and initial growth rate to those suitable for WT *Arabidopsis* RHs.

**Modelling WT**
To simulate the progression of a WT *Arabidopsis* RH through different growth stages into maturation, we adjust specific parameters from their baseline values (listed in Table S1) to reflect experimentally observed changes in tip growth rate (TGR) and cytoskeletal organization. Each simulation runs for 2000 time steps, corresponding to 1000 simulated minutes. The RH simulation begins with an initiation and growth stage lasting 800 time steps (400 min). Initially, the cell is spherical, representing the trichoblast. At $t = 0$, tip growth begins, and the RH starts to extend as a long, thin protrusion. When the RH grows long enough that the resting positions of the springs representing the cytoskeletal effects extend beyond the center of the initial spherical cell (around 7 $\mu m$ of tip extension), the nucleus forces are activated, causing the nucleus to start moving. All parameters are then kept constant during the remainder of the growth (G) phase. Following the G phase, we slow the tip growth rapidly over 100 time steps (50 min) to simulate the G/RG (reduced growth) transition. We then decreased TGR gradually over 300 time steps (150 min) in the RG phase before reaching maturation. Tip growth ceased after 1200 time steps (600 min), marking the transition to the early maturation (EM) phase, which continued for another 800 time steps (400 min).
The TGR is imposed within the model by varying the cell surface extensibility magnitude $\phi$. During the G/RG transition phase it decreases linearly to 0.2 of its original value over 100 time steps (50 min). It then decreases linearly to zero over the following 300 time steps (150 min) representing the RG phase to bring the tip growth rate to zero, where it remains during the EM phase.
To reflect the changes in the cytoskeleton intensity observed from experiments we also vary the strength of the microtubules $A_m$ into maturation. We decrease $A_m$ to 0.1 of its original value in the G/RG transition before decreasing it to zero over the following 300 time steps (150 min) of the RG phase, where it remains for the EM phase. At maturity the center of the actin potential $B_a$ increases linearly at a rate of approximately 0.3 units per time step, reflecting the experimentally observed retreat of the actin intensity peak.

**Sensitivity analysis of WT growing RHs**
Given the number of parameters in the mathematical model, we conducted a one-way sensitivity analysis to identify which parameters most influence the nucleus's morphology and position. The nucleus position relative to the tip is most influenced by $B_a$ , so actin effects have a greater impact on this metric compared to microtubule effects. This likely reflects the stronger influence of nucleus-associated forces compared to microtubule forces ($A_a > A_m$). Nucleus morphodynamic properties were primarily influenced by the same parameter of nucleus strength $\beta_{Nb} + \beta_{Nc}$, highlighting that these morphodynamic properties are not independent of each other. Changes in tip-nucleus speed, aspect ratio change speed, and area change speed are all most influenced by $A_a$. In contrast, circularity speed is most affected by the nucleus strength parameters $\beta_{Nb} + \beta_{Nc}$. For all four measures, the second most influential factor is the magnitude of noise $\alpha$.

**Modelling drug treatments**

Similar to modeling WT RH growth, to simulate Oryzalin and Latrunculin-B drug treatments, we adjust model parameters during the treatment and recovery phases. Reflecting experiments, the treatment phase takes 10 simulated mins (20 time steps) and the recovery phase is 90min long (180 time steps). To simulate treatment in G phase or RG phase, we only change the start time of treatment but relative changes in parameters are otherwise the same.

Under 10 min Oryzalin treatment, the microtubules depolymerise so we decrease the strength of the microtubule effects $A_m$to zero. TGR also decreases substantially, and sometimes tip bulging is observed. To simulate these effects, we reduce the extensibility magnitude $\phi$ to 10% of its original value $(0.1\phi)$ to slow the TGR while allowing the tip region size $L$ to increase to $1.3L$ enabling bulging. To avoid local biologically unrealistic behaviours such as temporary local increases in TGR, the time scales these changes occur over can be different but all reach their end of treatment value within 10 min. Here we set $A_m$ to drop to zero immediately; $L$ increases linearly over the first 5 min and $\phi$ decreases linearly over the full 10 min. During the 90-min recovery phase, $L$ linearly returns to its pre-treatment value eliminating bulging. However $\phi$ only partially recovers reaching 55% of its original value $(0.55\phi)$ to increase TGR but, as experimentally observed, not back to its pre-treatment level. The microtubule effect strength $A_m$ returns to its pre-treatment value but recovers in the last 30 mins of the 90-min recovery phase.

Under 10 min Latrunculin-B treatment, the actin depolymerises so we reduce the strength of the actin effects $A_a$ to zero. The TGR is also observed to drop to zero so we linearly reduce the cell extensibility magnitude $\phi$ to zero over the simulated 10 min treatment time. During the 90 min recovery phase, experimental observations show the TGR does not recover, so we retain the extensibility at 0. Actin filaments partially regenerate so we increase the strength of the actin effects but return it to only 10% of its original value $(0.1A_a)$ over this recovery period. To account for the crosstalk of AFs on MTs, we also allow $A_m$ to change at the same rate as the $A_a$ (decreasing linearly to zero and then partially recovering) but delayed by 20 min (although this effect is minimal). For parameters changes see Table S2.

**Sensitivity analysis of drug treatment parameters**

We perform a one-way sensitivity analysis to determine the influence of treatment specific parameters changes on the nucleus position and AR kinetic slope; nucleus position change over 21 min control followed by treatment into the start of recovery; and 21min at the end of recovery. With Oryzalin treatment time scales we observe that the MT recovery start time $MT_{recov}$ has the largest effect on nucleus position change in recovery whereas the depolymerization time $MT_1$ has the largest effect on the other measures. Excluding time scales, the increase in the size of the growth zone $L_{inc}$ effects the nucleus treatment position the most; $\phi_{recov}$ effects AR kinetic slope most; whereas the reduction in extensibility $\phi_{nuc}$ affects the two other measures the most. In Latrunculin-B treatments the recovery of actin during the recovery phase has the largest effect on the nucleus position change in treatment, which is larger than the Oryzalin parameters. Its effect on nucleus AR, while small, is also larger than the Oryzalin treatment parameters.

**Modelling mutants**

To model the *fra2* and *crwn1-2* mutants, we adjust parameters from their WT default values. In *fra2*, the TGR is observed to decrease sharply to zero. Therefore we set the extensibility magnitude $\phi$ at its WT G phase value until 25 min before tip growth termination. The

extensibility magnitude $\phi$ then decreases linearly over 25 min to zero to stop tip growth. The microtubule strength $A_m$ remains constant at its WT growth value and never decreases. The decrease in $\phi$ occurs either after 400 min to coincide with the RG stage or at 575 min to occur immediately before the start of the EM phase.
For *crwn1-2*, the TGR is observed to be slower. Therefore, we set the extensibility as $10\%$ of its WT value throughout the simulation. When we additionally perturb the nucleus parameters, we reduce nucleus volume to halve its WT value and reduce the nucleus rest length $\beta_N$ to half its WT value (by halving $\beta_{Nb}$ and $\beta_{Nc}$), while increasing the nucleus spring force $\alpha_N$ by a factor of 10.

**References**

[1] J. Dumais, S. L. Shaw, C. R. Steele, S. R. Long, P. M. Ray. An anisotropic-viscoplastic model of plant cell morphogenesis by tip growth. Int J Dev Biol. 2006;50(2-3):209-22.

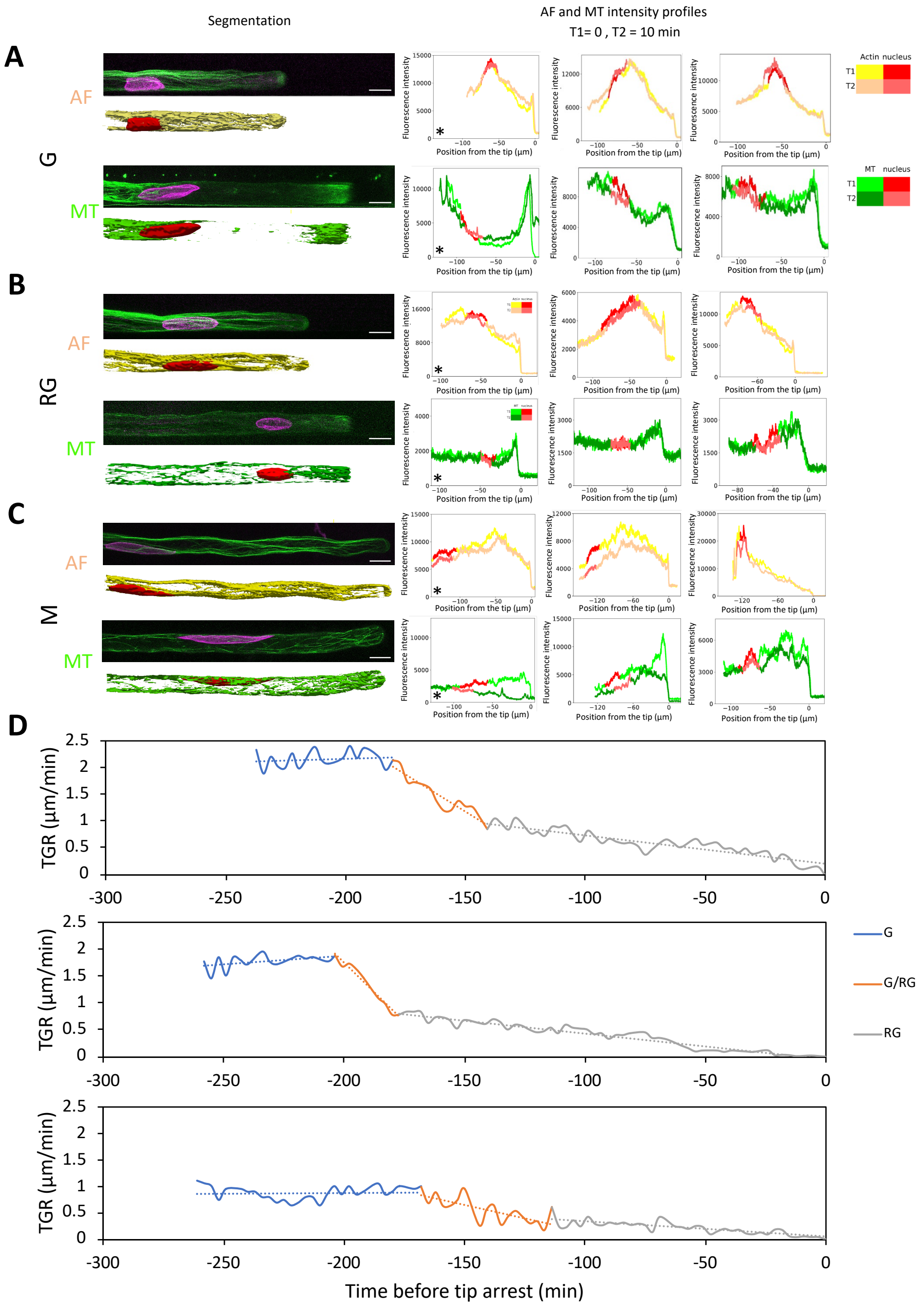

**Fig. S1: Cytoskeleton organization in various RHs during developmental stages and TGR kinetics. (A** to **C)** Confocal microscopy images and cumulated Microtubule (MT) and Actin filament (AF) fluorescence intensity of RHs from each stage: G (**A**), RG (**B**), and EM (**C**). White scale bar: 10µm. Star represents fluorescence profiles corresponding to the images on the left. (**D**) TGR kinetics for 3 individual RHs with fitting curves for G (blue) and RG (grey) growth phases as well as the G/RG transition (orange).

**A**

| | Nucleus-Tip Distance | AR | Circularity | Area | Nucleus-Tip Speed | AR Speed | Circularity Speed | Area Speed |
|---|---|---|---|---|---|---|---|---|
| $B_a$ | 0.76 | 0.016 | 0.012 | 0.00086 | 0.53 | 0.2 | 0.065 | 0.14 |
| $B_m$ | 0.1 | 0.16 | -0.13 | 0.052 | 0.94 | 0.8 | 0.79 | 0.8 |
| $A_m$ | 0.055 | 0.052 | 0.0058 | 0.0084 | 0.67 | 1.2 | 0.91 | 1.1 |
| $A_a$ | 0.33 | -0.087 | 0.017 | -0.019 | -1.9 | -1.7 | -1.3 | -1.5 |
| $\alpha$ | -0.16 | 0.013 | 0.019 | -0.00096 | 1.2 | 1.3 | 1.6 | 1.4 |
| $\alpha_N$ | -0.17 | 0.2 | -0.2 | 0.073 | -0.58 | 0.78 | 0.85 | 0.81 |
| $\beta_{Nb} + \beta_{Nc}$ | -0.15 | 2 | -1.7 | 0.68 | -0.65 | 0.22 | -2.3 | -0.93 |

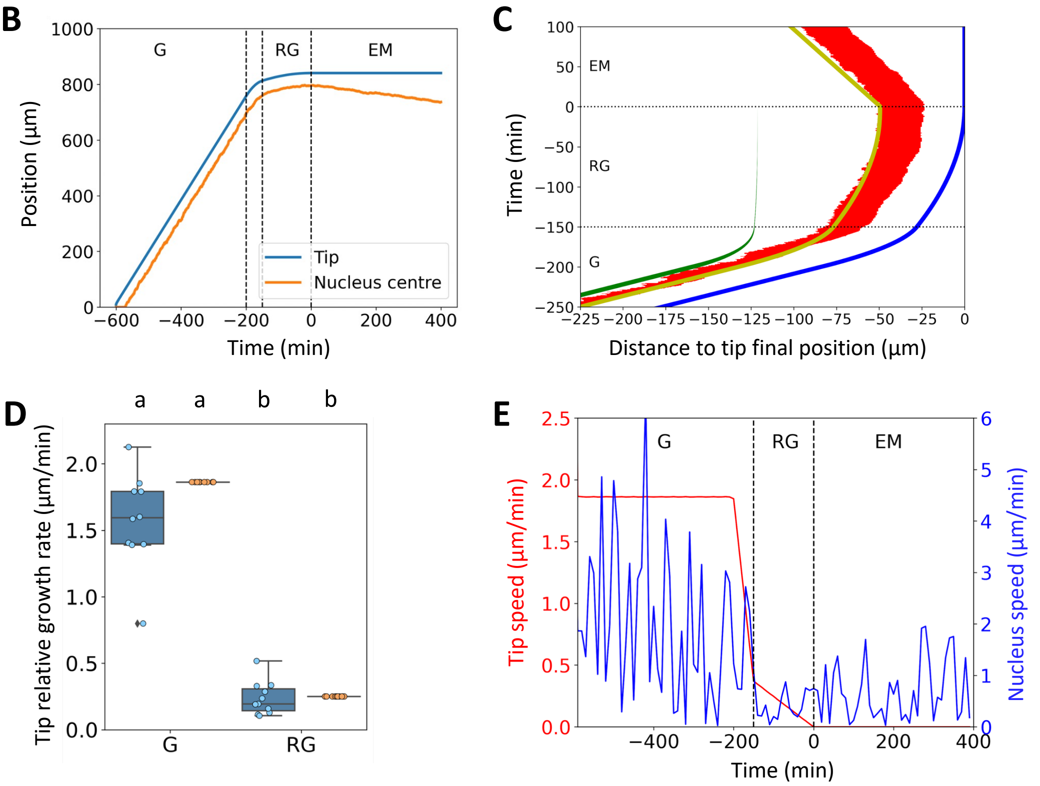

**Fig. S2: Supplementary information related to the model. (A)** Sensitivity analysis of nucleus parameters with maximum absolute values for each measure highlighted in yellow. (**B**) Simulation of the evolution tip (blue) and nuclear positions (orange) relative to growth initiation. (**C**) Simulation of nuclear position (red marking full width) relative to the tip (blue) with the central position of the MT (green) and AF (yellow) effects. (**D**) Comparison of TGR between G and RG stages showing both the model (orange) and experimental (blue) data. Different letters highlight statistically different conditions according to Mann-Whitney non parametric test ($p < 0.05$). (**E**) Simulation of RH tip speed (red) and nuclear speed (blue) over time.

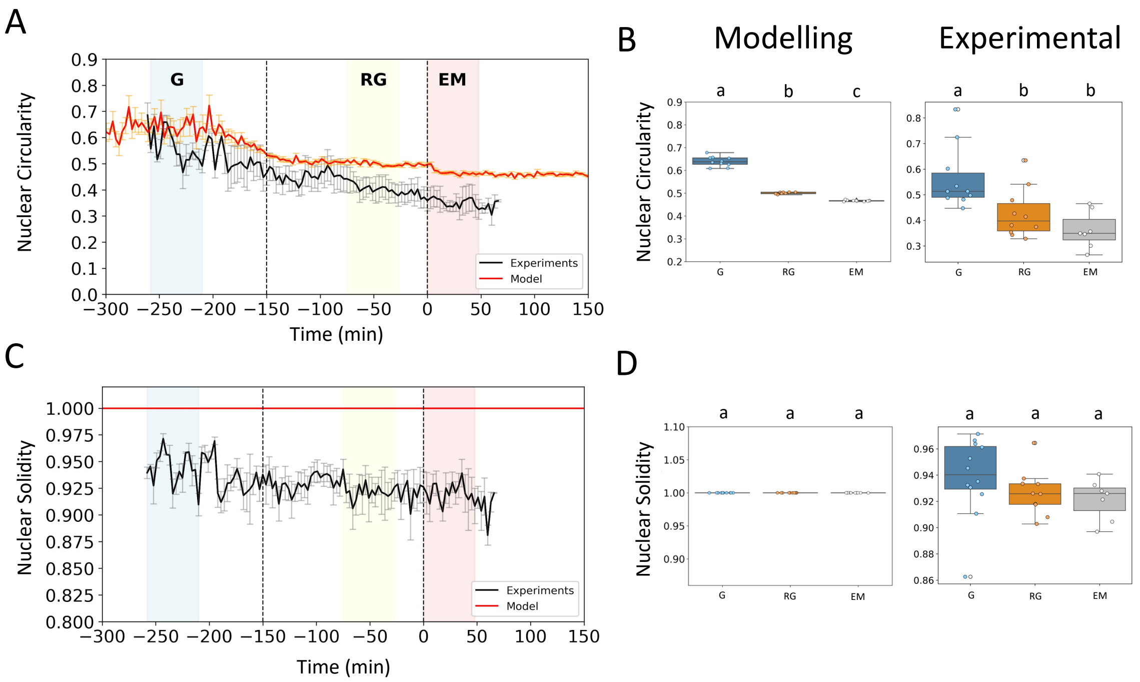

**Fig. S3: Additional nuclear dynamics parameters**. (**A**) Superposition of RH experimental kinetics (black line) and simulation data for nuclear circularity (**C**) and nuclear solidity. Blue, yellow and red boxes highlight the time frame used to produce average values for each growth phase. Dispersion of model (left) and experimental (right) data for nuclear circularity (**B**) and nuclear solidity (**D**) between each growth phases. Different letters highlight statistically different conditions according to Mann-Whitney non parametric test ($p < 0.05$).

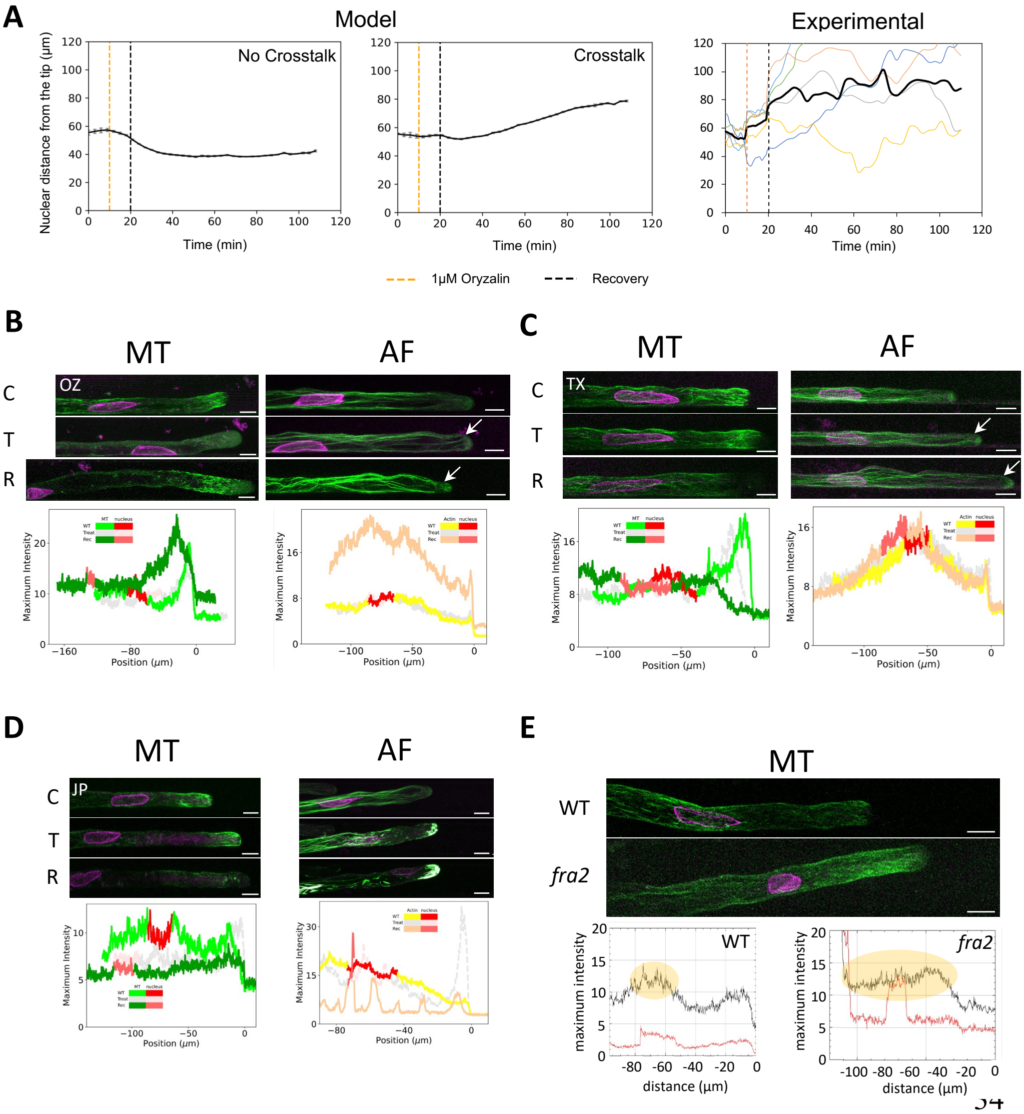

**Fig. S4: Simulation of AF-MT crosstalk and cytoskeleton network organization during drug treatment and in the *fra2* mutant. (A)** Simulation **of** nuclear movement upon 1 µM Oryzalin (OZ) treatment and after drug removal in recovery with $A_{ct}$ =1 (right) or without $A_{ct}$ =0 (left) presence of a AF-MT crosstalk. The panel on the far right shows the experimental kinetics of the different RHs used in the experiment (coloured lines) with the average (black line). (**B** to **D)** Confocal imaging of MT (left) and AF (right) organization of RHs treated with 1µM OZ (B), 10µM Taxol (C, White arrows: AF bundling progression in the subapical region) and 5µM Jasplakinolide (D). The cytoskeleton organization is compared between control (C), treated (T) and recovery (R) conditions. MT was tracked using GFP-MBD, AF using LifeAct-GFP and the nucleus with SUN2-tagRFP. The fluorescence profiles of both cytoskeleton across each condition is displayed underneath, with the nucleus position highlighted in red on each curve. **(E**) Confocal image of *fra2* mutant compared to WT Col-0. MTs were tracked using GFP-MBD and the nucleus with SUN2-tagRFP. The longitudinal fluorescence profiles for both channels (MTs in black and nucleus in red) are displayed below in relation to the tip position (0). White scale bar: 10µm

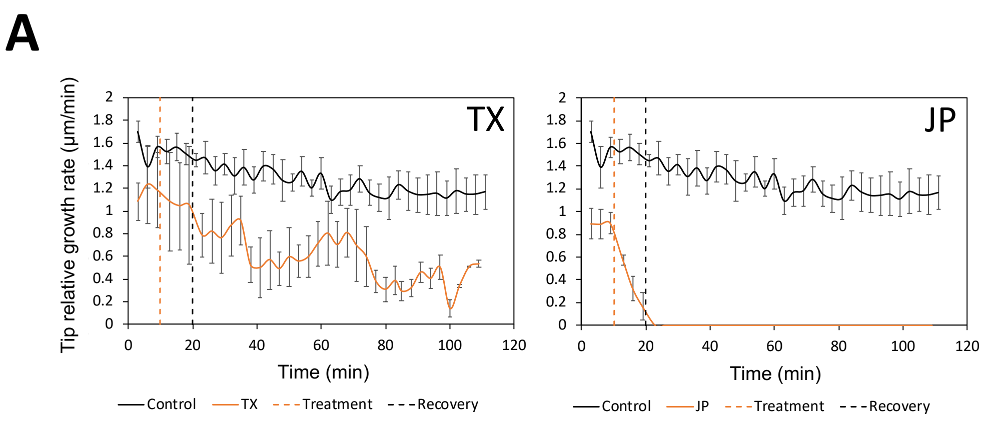

B

| | Nucleus position kinetic slope | Nucleus AR kinetic slope | Nucleus position treatment | Nucleus position recovery |
|---|---|---|---|---|
| $MT_1$ | -0.009 | 0.00021 | -0.42 | -0.32 |
| $MT_2$ | 0.00059 | 2.9e-05 | -0.17 | 0.21 |
| $A_1$ | 0.00083 | 2.4e-05 | -0.0095 | 0.094 |
| $MT_{recov}$ | 0.003 | 5.6e-05 | -0.0038 | 0.37 |
| $L_{inc}$ | 0.46 | -0.00057 | 6.9 | 26 |
| $\phi_{nuc}$ | -1.7 | 0.0049 | -1 | -1.4e+02 |
| $\phi_{recov}$ | 0.64 | -0.011 | 6.1 | 66 |
| $LatA_{recov}$ | -0.29 | 0.029 | -1.2e+02 | -6.8 |

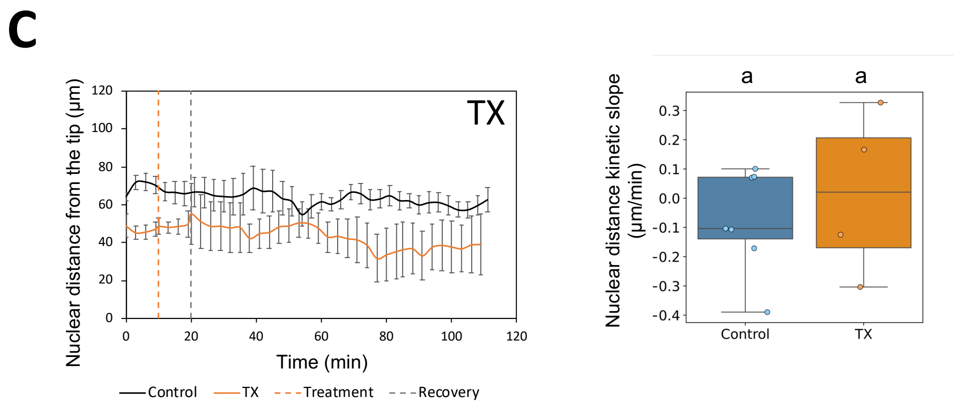

**Fig. S5: Sensitivity analysis of treatment parameters**. (**A**) Kinetics of TGR of G phase RH in mock conditions (black line) or treatments with either 10µM Taxol (TX) or 5µM Jasplakinolide (JP). The kinetics correspond to a 10 min-control medium, followed by a 10 min-treatment, and a 90 min-drug removal. (**B**) Sensitivity analysis of nucleus parameters during treatment and recovery with maximum absolute values for each measure highlighted in yellow. (**C**) Kinetics of nuclear distance from the tip in G phase RH during control conditions (black line) or treated with 10µM Taxol (TX). The slope of both kinetics is displayed as boxplots on the right.

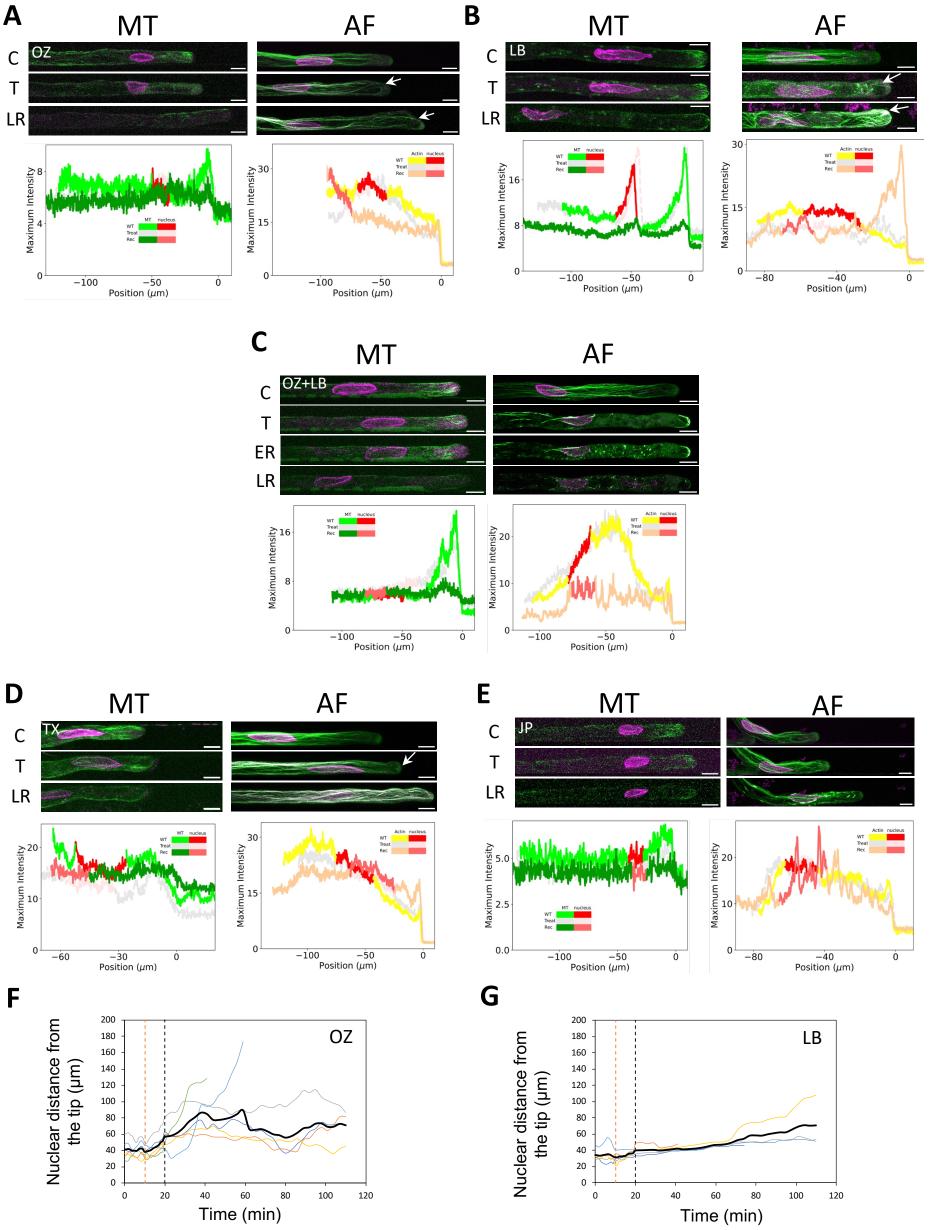

**Fig. S6: MT and AF network organization in RG phase during cytoskeleton targeting drug treatment. (A to G). A to E**: Confocal imaging of MT (left) and AF (right) organization of RG phase RHs treated with 1µM Oryzalin (OZ), (**A**), 1µM Latrunculin B (LB), (**B**), both 1µM Oryzalin and 1µM Latrunculin B (**C**), 10µM Taxol (TX) (**D**) and 5µM Jasplakinolide (JP) (**E**). The cytoskeleton organization is compared between control (C), treated (T) and late recovery (LR) conditions in OZ and LB treatments. In the double treatment with both OZ and LB, the early recovery (ER) is indicated to show the delay in the response to the treatment. MTs were tracked using GFP-MBD, AFs using LifeAct-GFP and the nucleus position with SUN2-tagRFP. White scale bar: 10µm. The fluorescence intensity profiles for both cytoskeleton across each condition is displayed underneath. The nucleus position is highlighted in red on each curve. Nuclear distance kinetics upon 1µM Oryzalin (**F**) or 1µM Latrunculin B (**G**) treatment and recovery in individual RG RHs (coloured lines) and in average (bold black line). The orange dashed line indicates the start of the treatment and the black one the end of treatment.

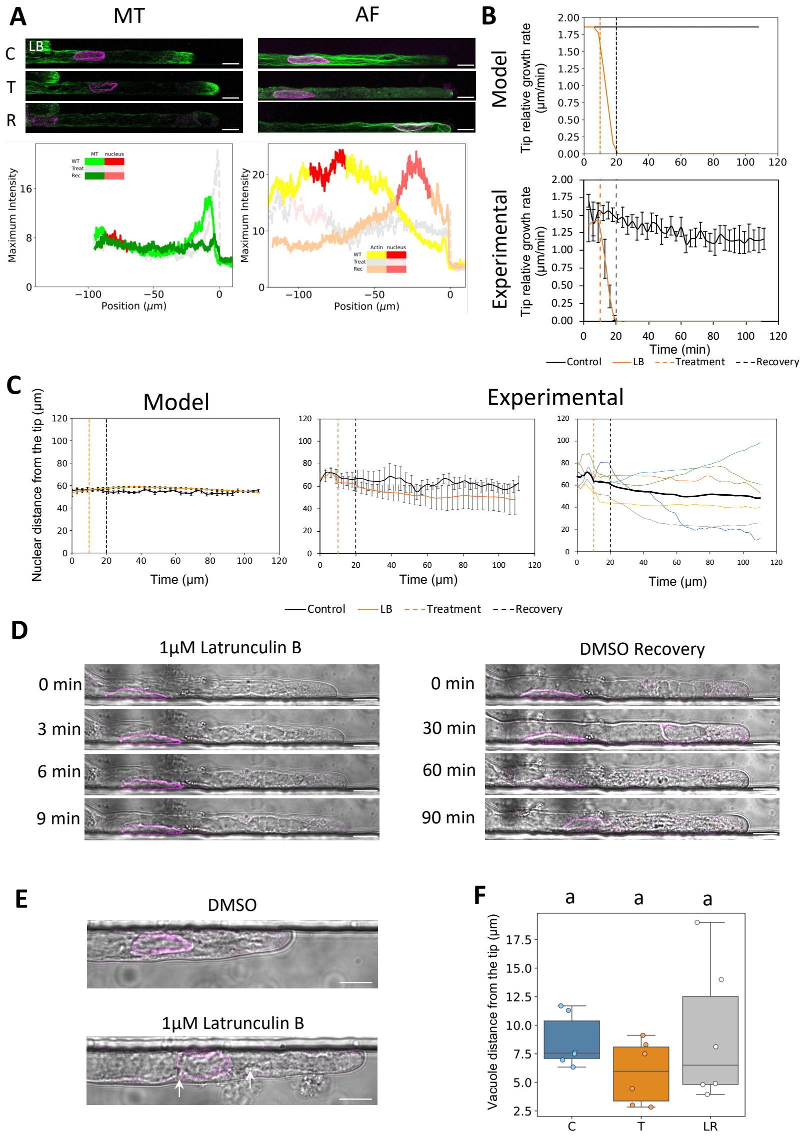

**Fig. S7: MT and AF network organization in G phase upon AF destabilization and recovery. (A)** Confocal imaging of MT (left) and AF (right) organization of G phase RHs treated with 1µM Latrunculin B (LB). The cytoskeleton organization is compared between control (C), treated (T) and late recovery (LR) conditions. MT was tracked using GFP-MBD, AF using LifeAct-GFP and the nucleus with SUN2-tagRFP. White scale bar: 10µm. The fluorescence intensity profiles for both cytoskeleton across each condition is displayed underneath. The nucleus position is highlighted in red on each curve. (**B**) Kinetics of TGR of G phase RHs in mock conditions (black line) or treatments with 1µM LB (orange line) from both model simulations (above) and experimental data (below). The kinetics correspond to a 10 min-control medium, followed by a 10 min-treatment, and a 90 min-drug removal. (**C**) Kinetics of nuclear distance from the tip of G phase RHs in control conditions (black line) or treated with 1µM LB (orange line) in simulations (left) and experiments (right). The far-right panel shows individual RHs kinetics (coloured lines) and the average kinetics (black line). (**D**) Confocal DIC images showing the evolution of vacuole fragmentation in G phase RHs upon 10 min LB treatment and 90 min of drug-removal recovery. The nucleus is highlighted with SUN2-tagRFP. (**E**) Confocal DIC images showing the evolution of vacuole fragmentation in G and RG phases RHs around the nucleus upon LB treatment compared to Control. White scale bar: 10µm. (**F**) Vacuole position in G phase RHs during control (C), treated (T) and late recovery (LR) conditions. Different letters highlight statistically different conditions according to Mann-Whitney non parametric test ($p < 0.05$).

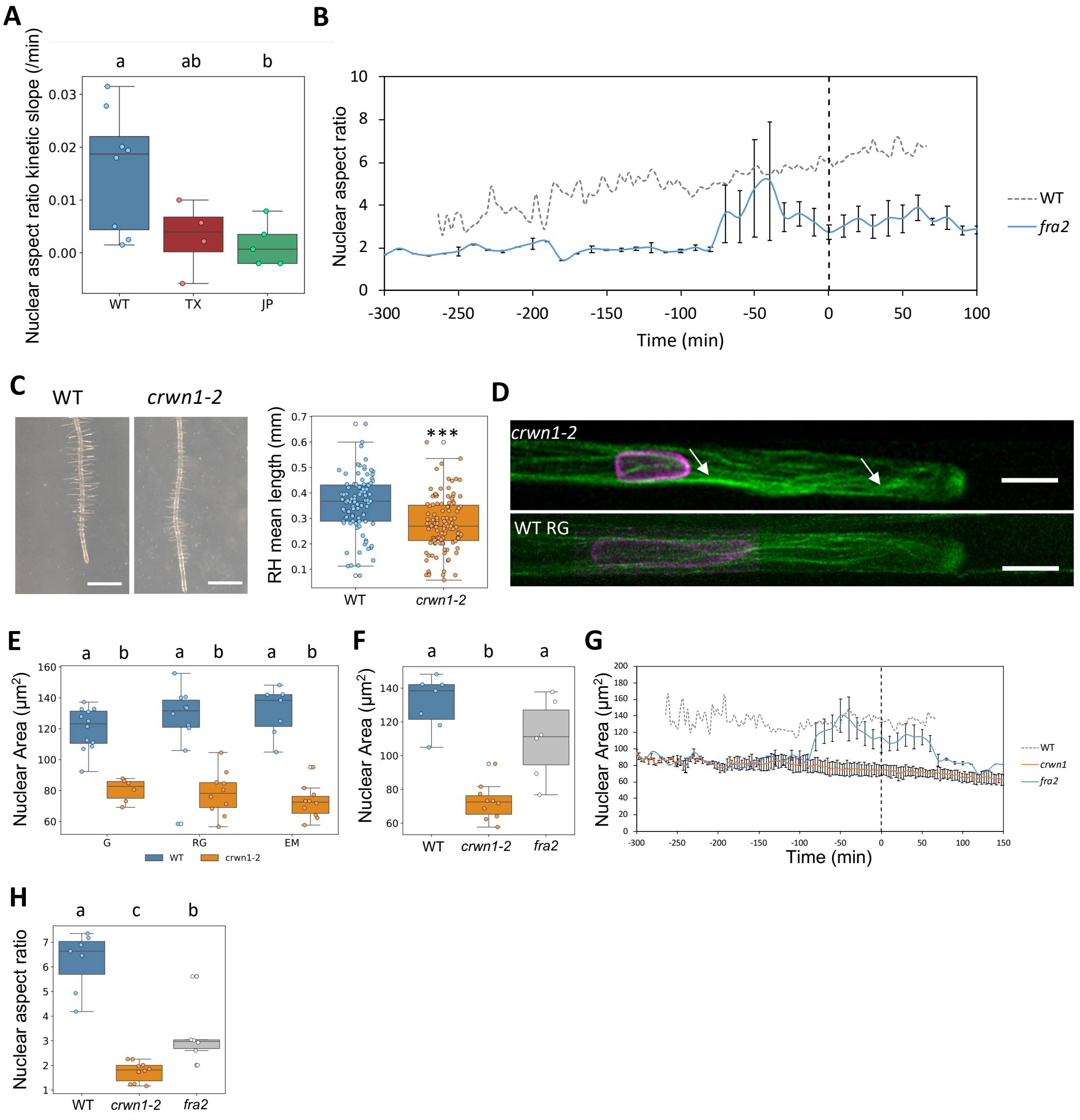

**Fig. S8**: **Nuclear morphology after AF or MT stabilization or nucleoskeleton mutation.** **(A)** Nuclear AR kinetic slope derived from kinetics of TX and JP treatment and recovery in RG phase RHs. (**B**) Kinetics of *fra2* nuclear aspect ratio (blue line) in comparison to WT (dashed grey line). (**C**) Mean RH length in 9-DAG seedlings from WT and *crwn1-2* mutant lines. ***: p-value <0.001. (**D**) Confocal images of *crwn1-2* and WT Col-0 RH with AFs (LifeAct-GFP) and nuclear envelope (SUN2-tagRFP). RHs from *crwn1-2* mutants show an increase in AF bundling (white arrows). White scale bar: 10µm. (**E**) Nuclear area between Col-0 and *crwn1-2* throughout the different growth stages. (**F**) Nuclear AR after tip arrest in *fra2* in comparison to WT Col-0 and *crwn1-2*. (**G**) Kinetics of nuclear area in *crwn1-2* (orange line) and *fra2* (blue line) in comparison to WT Col-0 (dashed line). (**H**) Nuclear AR in *fra2* and *crwn1-2* compared to WT after tip growth arrest. Different letters highlight statistically different conditions according to Mann-Whitney non parametric test ($p < 0.05$).

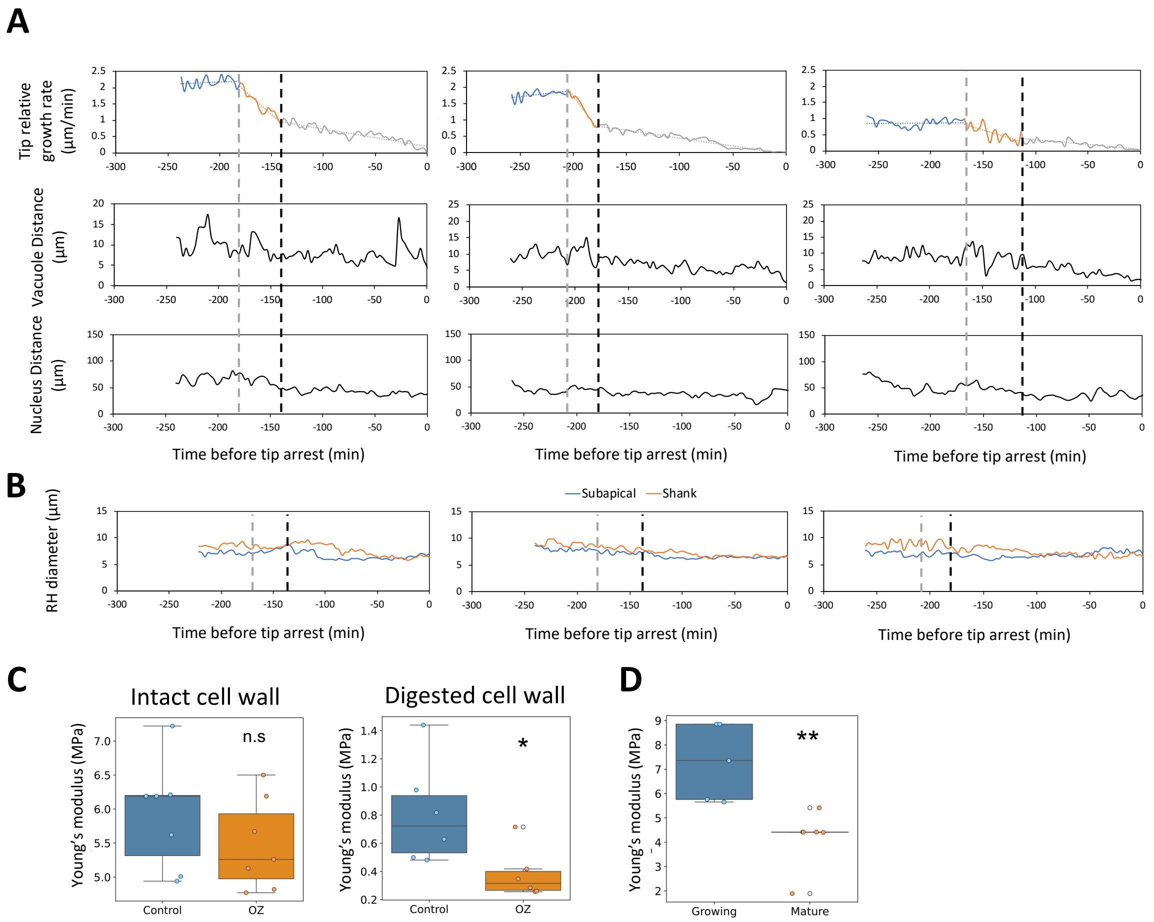

**Figure S9: Synchronicity of RH growth parameters changes during maturation:** (**A**) Synchronisation of vacuole and nuclear distance variations with the transition between G and EG from the tip in three RH examples. (**B**) Evolution of RH diameter in shank (orange line) and in subapical region (blue line) from the tip in three RH examples. (**C**) Effect of 1µM Oryzalin treatment on cell stiffness (C) with an intact RH (left) and with an enzymatically semi-digested cell wall (right) *: p value <0.05. (**D**) Cell stiffness between growing and mature RHs. **: p value <0.01

**Table S1**: List of parameters used in the mathematical model for WT RHs during growth.

| Parameter | Symbol | Value |
|---|---|---|
| Tip growth region length | $L$ | 1.57 |
| Extensibility magnitude initially | $\phi$ | 1.4 |
| Extensibility shape power | $b$ | 10 |
| Poison ratio | $\nu$ | 0.5 |
| Pressure | $P$ | 5 |
| Cell wall thickness | $\delta$ | 1 |
| [redacted] | | |
| Microtubule strength initially | $\tilde{A}_m$ | 0.005 |
| Actin strength initially | $\tilde{A}_a$ | 0.05 |
| Microtubule position | $B_m$ | 8 |
| Actin position | $B_a$ | 5 |
| Nucleus spring force | $\alpha_N$ | 0.2 |
| Nucleus spring length actin independent | $\beta_{Nb}$ | 1.3 |
| Nucleus spring length actin dependent | $\beta_{Nc}$ | 1.7 |
| Cross-talk magnitude | $A_{ct}$ | 1 |
| Cross-talk scaling | $C_{ct}$ | 100 |
| Cross-talk scaling power | $C_{ctf}$ | 1 |
| Noise standard-deviation (SD) magnitude | $\alpha$ | 30 |
| Noise SD actin independent proportion | $\beta$ | 0.05 |
| Noise SD microtubule independent proportion | $\gamma$ | 0.5 |
| Boundary force interaction magnitude | $\zeta$ | 100 |
| Boundary force interaction distance | $\eta$ | 0.1 |
| Boundary force nucleus width magnitude | $\zeta_N$ | 10000 |
| Boundary force nucleus minimum length | $\eta_N$ | 1 |
| Nucleus volume | $V$ | 0.15 |

**Table S2:** Parameter changes during Oryzalin and Latrunculin-B drugs treatments and recovery afterwards which are used for the sensitivity analysis of Fig S5B.

| Parameter | Symbol | Value |
|---|---|---|
| Nucleus end MT depolymerization time | $MT_1$ | 400 |
| Tip MT depolymerization time | $MT_2$ | 410 |
| Actin depolymerisation time | $A_1$ | 420 |
| Time MT recovery begins | $MT_{recov}$ | 540 |
| [redacted] | | |
| L increase during OZ treatment | $L_{inc}$ | 0.3 |
| Reduction in tip extensibility | $\phi_{nuc}$ | 0.9 |
| Level of extensibility recovery | $\phi_{recov}$ | 0.5 |
| [redacted] | | |
| Actin recovery during LB treatment | $LatA_{recov}$ | 0.1 |
